# NicheFlow: Towards a foundation model for Species Distribution Modelling

**DOI:** 10.1101/2024.10.15.618541

**Authors:** Russell Dinnage

## Abstract

1. Species distribution models (SDMs) are crucial tools for understanding and predicting biodiversity patterns, yet they often struggle with limited data, biased sampling, and complex species-environment relationships. Here I present NicheFlow, a novel foundation model for SDMs that leverages generative AI to address these challenges and advance our ability to model and predict species distributions across taxa and environments.

2. NicheFlow employs a two-stage generative approach, combining species embeddings with two chained generative models, one to generate a distribution in environmental space, and a second to generate a distribution in geographic space. This architecture allows for the sharing of information across species and captures complex, non-linear relationships in environmental space. I trained NicheFlow on a comprehensive dataset of reptile distributions and evaluated its performance using both standard SDM metrics and zero-shot prediction tasks.

3. NicheFlow demonstrates good predictive performance, particularly for rare and data-deficient species. The model successfully generated plausible distributions for species not seen during training, showcasing its potential for zero-shot prediction. The learned species embeddings captured meaningful ecological information, revealing patterns in niche structure across taxa, latitude and range sizes.

4. As a proof-of-principle foundation model, NicheFlow represents a significant advance in species distribution modeling, offering a powerful tool for addressing pressing questions in ecology, evolution, and conservation biology. Its ability to model joint species distributions and generate hypothetical niches opens new avenues for exploring ecological and evolutionary questions, including ancestral niche reconstruction and community assembly processes. This approach has the potential to transform our understanding of biodiversity patterns and improve our capacity to predict and manage species distributions in the face of global change.

## Introduction

The accelerating pace of environmental change has amplified the need for accurate species distribution predictions, a cornerstone of biodiversity conservation, ecological research, and informed management decisions. Species distribution models (SDMs) have become indispensable tools for mapping and forecasting species occurrences under current and future conditions, playing a crucial role in efforts to mitigate the impacts of habitat loss, climate change, and other anthropogenic pressures. However, traditional SDMs often stumble when confronted with rare or data-deficient species, typically demanding substantial occurrence data and leading to repetitive, species-specific modeling efforts across research groups and conservation practitioners (Guisan et al., 2017).

Conventional SDMs, such as Maxent, Generalized Linear Models (GLMs), and Random Forests (RF), rely heavily on species-specific occurrence records and environmental variables to estimate species-environment relationships (Elith & Leathwick, 2009). These models often operate under the assumption that species niches are determined solely by current environmental conditions, aligning with the environmental niche concept that defines a species’ fundamental ecological space based on its abiotic and biotic requirements (Sobeŕon, 2007). However, the variability in availability and quality of occurrence data can lead to biased or incomplete predictions, particularly for rare, cryptic, or newly discovered species (Yackulic et al., 2013).

The emergence of foundation models in ecology, particularly those leveraging generative AI approaches, offers a paradigm shift: a unified model capable of generating distribution predictions for hundreds of thousands of species, including those absent from its training data. This approach not only streamlines the modeling process but also unlocks the potential for robust predictions in the face of limited data, a common challenge in biodiversity research (Beery et al 2021).

### Unlocking the potential of foundation models in Ecology and Conservation

The potential of foundation models in ecology extends far beyond mere prediction. These models, which have revolutionized fields like natural language processing and computer vision (Bommasani et al., 2021), offer a suite of advantages that could transform ecological research:

1. Reduction of Duplicated Effort: A unified foundation model allows movement beyond the fragmented landscape of species-specific models, enabling collective progress and ensuring consistency across predictions (Pimm et al., 2015; Franklin, 2013).
2. Computational Efficiency: Pre-trained models significantly reduce the computational demands of SDMs, an increasingly important consideration given the rising concerns over the carbon footprint of machine learning (Strubell et al., 2019; Patterson & Hennessy, 2021).
3. Democratization of Advanced Techniques: By simplifying the modeling process, foundation models can make sophisticated analytical tools accessible to ecologists with limited machine learning expertise, broadening the community of researchers who can contribute to and benefit from cutting-edge SDM techniques (Beery et al. 2021).
4. Collaborative Model Improvement: A unified model fosters a cycle of iterative improvement, where each user builds upon the work of others, enhancing model performance over time (Pereira et al., 2010).

In this paper, I present a novel approach that combines generative AI with species embeddings derived from distribution data, enabling zero-shot predictions in SDMs without requiring explicit trait or phylogenetic information. This method builds upon recent advances in machine learning, particularly in the fields of generative modeling and representation learning (Reichstein et al., 2019; Ho et al., 2020). By learning latent representations of ecological niches, the model aligns with the concept that niches are defined by where a species occurs relative to environmental gradients (Sobeŕon, 2007).

### Enabling Zero-shot Species Distribution Prediction

An exciting aspect of foundation models is their capacity for few-shot and zero-shot learning. Few-shot learning refers to the ability of a model to make accurate predictions with very limited training data for a particular task or category (Wang et al., 2020). Zero-shot learning goes a step further, allowing models to make predictions for entirely new categories that were not present at all in the training data (Xian et al., 2018). These concepts, while originating from machine learning, have profound implications for ecology. In the context of SDMs, zero-shot learning would enable predictions of species distributions for which we have no occurrence data in our training set. This capability is analogous to an experienced ecologist making an educated guess about where a newly discovered species might occur based on its taxonomic relationships and the known distributions of similar species (Lampert et al., 2014). For SDMs, this means we could potentially predict distributions for rare, newly discovered, or data-deficient species by leveraging the model’s learned representations of ecological niches and species-environment relationships across a wide range of taxa (Norberg et al., 2019).

### Capturing ecological meaning

The model I present here goes beyond traditional Joint Species Distribution Models (JSDMs) by capturing the “distribution of distributions that is, the underlying environmental niches of species. Instead of focusing on the residual covariance among species, as in linear JSDMs (Pollock et al., 2014; Ovaskainen et al., 2016), this generative AI approach seeks to learn the distribution of ecological niches directly from occurrence data. This allows for the estimation of complex, multidimensional patterns that define species’ environmental tolerances with a flexibility and power that surpasses traditional JSDMs (Norberg et al., 2019).

Beyond its predictive applications, the species embeddings generated by this model serve as a powerful research tool, encoding ecological niches in a way that allows for downstream analyses such as estimating ecological distances between species, reconstructing ancestral niches, or querying for species with similar environmental tolerances. This functionality provides a unique opportunity to explore the ecological dimensions of biodiversity, deepening our understanding of species’ fundamental and realized niches and their evolutionary implications (Sobeŕon & Peterson, 2005).

In the following sections, I detail the technical implementation of this approach, present results demonstrating its performance on both seen and unseen species, and discuss the broader implications for ecological research and biodiversity conservation. This work represents a significant step towards unifying species distribution knowledge into a single, powerful predictive framework, opening new avenues for addressing pressing ecological challenges and advancing our understanding of biodiversity patterns and processes.

## Methods

### Generative Model Framework for Joint Species Distribution Modeling

I develop a generative approach to species distribution modeling that integrates species-specific information and environmental conditions through probabilistic models. This approach allows us to capture complex ecological relationships by linking species occurrences with environmental variables and geographic coordinates.

### Model Equation

In order to create a usable generative model for species distribution modeling it needs to have a target probability distribution to generate from. To create a map of species the goal is to sample from the probability distribution of species across geographic coordinates (*X, Y*), given that the species is *S* = *s*, and that it occurs (*O_s_* = 1) e.g. *P* (*X, Y |S* = *s, O_s_* = 1). We further need to condition on species in a quantitative generalizable way. To do this, instead of conditioning on the identity of a species, we can condition on a vector representation of the species’s niche, a latent vector-valued variable we will call *Z*, which will be estimated by the model along with the other parameters. For simplicity I will use the expression *Z* = *z_s_* to represent *S* = *s, O_s_* = 1, leaving us with *P* (*X, Y |Z* = *Z_s_*). To include this latent niche vector and also the environment in our probability of interest, we can use the law of total probability to arrive at the following mathematical representation:

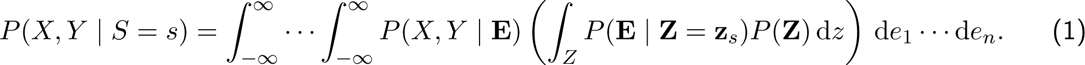

This equation shows how the occurrence of a species can be modeled by chaining two probability distributions: one that describes the species’ environmental niche (*P* (**E** *| Z* = *z_s_*)) and another that links environmental conditions to geographic space (*P* (*X, Y |* **E**)). Figure 1 shows this two-step sampling process conceptually. By learning a representation of the species niche as a latent variable **Z***_s_*, we can create a flexible model that captures both the environmental dependencies and geographic patterns of species distributions. The distribution of *z* represents the ‘distribution of distributions’. More specifically, the distribution across species of edistributions in environmental and geographic space. See the supporting information for the full derivation of equation 1.

**Figure 1:**
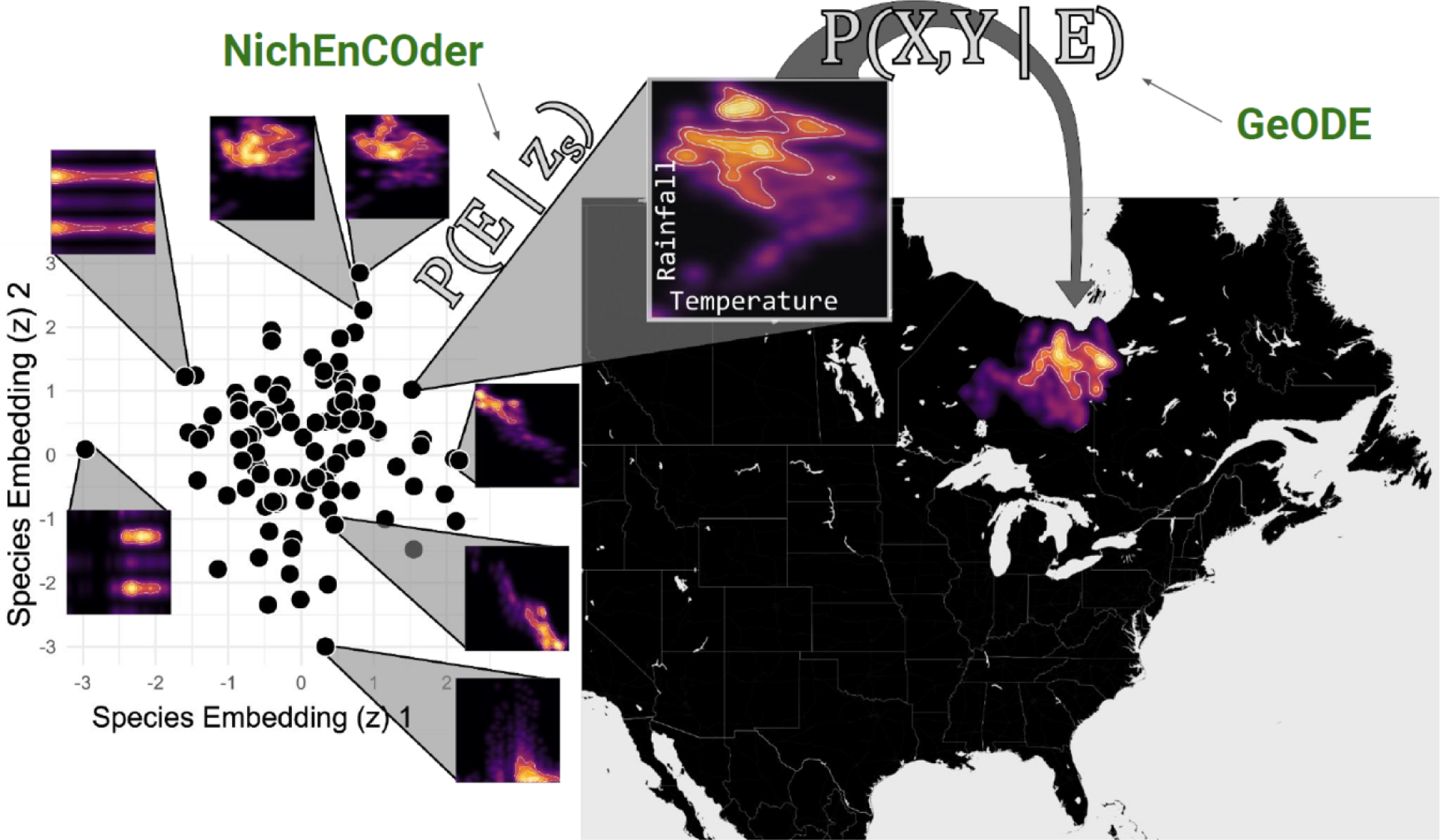
Conceptual illustration of the two step generative AI model for Joint Species Distribution Modeling called NicheFlow, proposed in this study. A generative model of the species environmental niche (NichEncoder) is composed with a generative model mapping environmental variables to geographic coordinates (GeODE). (GeODE). A species environmental niche is represented by a d-dimensional vector z that is transformed into a k-dimensional environmental probability distribution or hypervolume. A z vector for every species is estimated during model training and provides a generalizable, reusable compact representation of species’s niches. Across all species the distribution of z represents the ‘distribution of (environmental) distributions’.

### Sampling from the Species Distribution Using Generative Models

The equations derived above describe the probability of species occurrences as complex high-dimensional integrals that are computationally expensive to evaluate directly. To overcome this challenge, I leverage generative models, which can efficiently approximate these distributions by sampling, thus bypassing the need to compute these integrals explicitly.

### Generative Model Framework for Sampling

Generative models, such as Variational Autoencoders (VAEs) and rectified flow models, provide a powerful framework for approximating high-dimensional probability distributions through sampling. These models learn to generate data that resemble the distribution of the observed data by learning the underlying data-generating process.

## 1. Sampling from the Environmental Niche

From Equation (3), the term *P* (**E** *|* **Z** = **z***_s_*) represents the species’ environmental niche. We can use a generative model to learn this niche distribution by training it on environmental data associated with species occurrences. Once trained, the model can generate samples of environmental vectors **E** conditioned on the species embedding **Z** = **z***_s_*.

### Model Training

I train a generative model called **NichEncoder**, using a two-stage generative model – combination of a Condition Variation Autoencoder (CVAE; Zheng et al. 2023) and a Rectified Flow model (Liu et al. 2023). The model is trained using environmental occurrence data for many species. The model learns a mapping from a latent space (representing **Z**) to the environmental conditions that define species niches. See ‘Model Details’ for more details of NichEncoder.

### Sampling

After training, new environmental vectors **E** can be generated by sampling from the learned latent space. These samples represent possible environmental conditions under which the species *S* = *s* can occur.

## 2. Sampling Geographic Coordinates Given Environmental Conditions

The next step involves generating geographic coordinates (*X, Y*) given the sampled environmental conditions **E**. The term *P* (*X, Y |* **E**) describes this relationship and can also be modeled using a generative approach.

### Training the Spatial Generative Model

A second generative model is trained to map environmental conditions **E** to geographic coordinates. This model learns the spatial patterns of species occurrences based on the environmental vectors generated in the previous step. The model is called **GeODE**, and is based on a Conditional Rectified Flow model (Liu et al 2022). The name is based on the fact that Rectified Flow models estimate an Ordinary Differential Equation to transform noise into a complex high dimensional distribution. More details on **GeODE** can be found in the ‘Model Details’ section.

### Sequential Sampling

Once the spatial model is trained, it can sequentially generate coordinates (*X, Y*) by conditioning on the sampled environmental vectors. This process allows us to reconstruct the spatial distribution of the species without needing to evaluate the full integral.

## 3. Combining the Sampling Steps

By chaining the two generative models, the overall sampling process approximates the species distribution defined by Equation (3). Specifically:

- First, sample **E** from the environmental niche model conditioned on **Z** = **z***_s_*.
- Then, use the sampled **E** to generate corresponding coordinates (*X, Y*) using the spatial generative model.

This approach provides a flexible and efficient method to approximate the distribution of species occurrences, leveraging the generative model’s capacity to learn complex, high-dimensional relationships between species, environment, and geography.

### Zero-shot Species Distribution Modeling

Zero-shot Species Distribution Modeling (0-SDM) enables the estimation of geographic distributions for species not included in the training set by optimizing a latent embedding specific to the new species. This approach adjusts the embedding based on observed occurrence data by comparing predicted and observed environmental vectors using Energy Distance and Sinkhorn Distance. These distance measures provide a robust method for aligning predicted species distributions with observed data.

### Embedding Optimization

For a new species *S* = *s^∗^* not present in the training data, the goal is to find an optimal embedding **z***_s_∗* within the latent space learned by the generative models. This embedding is iteratively adjusted to fit the observed environmental conditions of the new species.

Steps of the Optimization Process:

## 1. Initialization

The species embedding **z***_s_∗* is initialized either randomly from the prior distribution *P* (**Z**) or based on similarities to embeddings of known species with similar ecological traits.

## 2. Sampling Environmental Conditions

The environmental generative model, defined as the function *f*_env_, is used to sample predicted environmental vectors 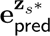 conditioned on the embedding **z***_s_∗*:

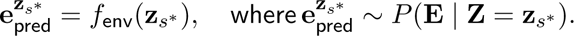

Here, *f*_env_ maps the species embedding to predicted environmental conditions, capturing the species’ ecological niche in the environmental space.

## 3. Loss Calculation Using Energy Distance and Sinkhorn Distance

To optimize the embedding **z***_s_∗*, we define a loss function that evaluates how well the predicted environmental vectors **E**_pred_ align with the observed environmental vectors **E**_true_ from occurrence data. Here, **E**_pred_ and **E**_true_ are matrices where rows represent individual environmental vectors associated with predicted and observed occurrences, respectively. These matrices can have different numbers of rows, reflecting the flexibility of the distance measures used. The overall loss function used for optimization is defined as:

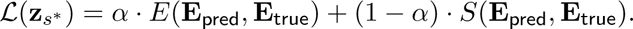

where *E* is Energy Distance (Sźekely & Rizzo, 2013) and *S* is the Sinkhorn Distance (Cuturi, 2013). Both are metric designed to estimated the similarity of two distribution expressed as point clouds. Using a combination of both balances their different strengths. See Supporting Information for details on Energy and Sinkhorn Distance including their equations, and optimization details.

## 4. Optimization of the Species Embedding

The species embedding **z***_s_∗* is optimized using stochastic gradient descent (SGD) to minimize the combined loss *L*(**z***_s_∗*). The iterative updates refine the embedding until the generated environmental predictions closely match the observed environmental data. #### Optimization Update:

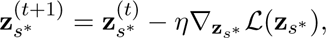

where *η* is the learning rate, and *∇***_z_***_s∗_ L* is the gradient of the loss function with respect to the embedding.

### NichEncoder: Generative Model for Species Environmental Niches

NichEncoder is a two-stage generative model designed to estimate species-specific environmental niches. It takes as input a vector of latent species embeddings, **z**_species_, and generates environmental variables, **e**, representing the conditions associated with species occurrences. This approach allows the model to learn complex, non-linear relationships between species and their environmental contexts, facilitating predictions of species distributions in novel scenarios. The model was implemented in R using the torch package, which provides a high-level interface to the PyTorch deep learning library.

### Model Architecture and Training

NichEncoder follows a two-stage generative approach inspired by the Two-Stage VAE architecture (Dai and Wipf, 2019), which is particularly useful for modeling complex, high-dimensional data distributions with structured priors. This architecture allows for dimensionality reduction and disentanglement of the latent space, improving the model’s ability to capture the underlying data manifold. In the context of NichEncoder, the first stage estimates the data manifold, and the second stage estimates the distribution of the data on this manifold. For the first stage I used a conditional Variational Autoencoder (CVAE: Zheng et. al 2023), and for the second stage I used a conditional Rectified Flow model (RF: Liu et al. 2023), where the generative models are both conditioned on *z_s_*, an estimated species-level latent niche variable. Details of the architectures can be found in the Supporting Information.

### Model Training and Implementation

Both stages of NichEncoder are trained sequentially using GPU acceleration with CUDA, with extensive logging and periodic checkpointing to monitor training progress and performance. The implementation of both stages was carried out in R using the torch package, which interfaces with the PyTorch library, allowing efficient and flexible model training in a high-level language environment.

### GeODE: Generative Model for Geographic Distributions

GeODE (Geographic Occurrence Distribution Estimator) is a generative model that employs a conditional rectified flow to predict geographic distributions (Figure 1). The model outputs longitude (*X*) and latitude (*Y*) coordinates by evolving a 2-dimensional noise vector toward the target distribution, which represents species’ occurrence points on the Earth’s surface. The transformation is guided by an Ordinary Differential Equation (ODE), which is conditioned on environmental vectors (**e**) corresponding to each *X, Y* pair. Unlike the NichEncoder model, GeODE does not require an initial VAE step because the output coordinates are already in a low-dimensional (2D) space, making the process more direct.

### Model Architecture

GeODE uses a modified rectified flow architecture similar to that used in NichEncoder but tailored specifically for geographic data. The model generates 2-dimensional noise vectors as inputs, which are transformed through the rectified flow mechanism to output the desired geographic coordinates. The input consists of random noise vectors representing initial guesses in 2D space, while the conditioning input (**e**) comprises environmental variables associated with each geographic location. Each environmental vector is normalized using means and standard deviations calculated from the data, ensuring numerical stability during training.

### U-net Architecture

The core of GeODE is a U-net style structure implemented with Multi-Layer Perceptrons (MLPs) instead of convolutional layers (Figure 2). The U-net consists of two primary paths: downsampling and upsampling. In the downsampling path, the input noise vectors and environmental conditioning are passed through three fully connected layers with progressively smaller neuron counts (512, 256, and 128). These layers reduce the dimensionality while learning broad, high-level representations of the relationship between geographic locations and environmental factors. The upsampling path reconstructs the geographic coordinates by reversing the dimensionality reduction, using three corresponding fully connected layers to produce the final outputs. Skip connections between the downsampling and upsampling paths retain and propagate finer details, leading to more accurate predictions.

**Figure 2:**
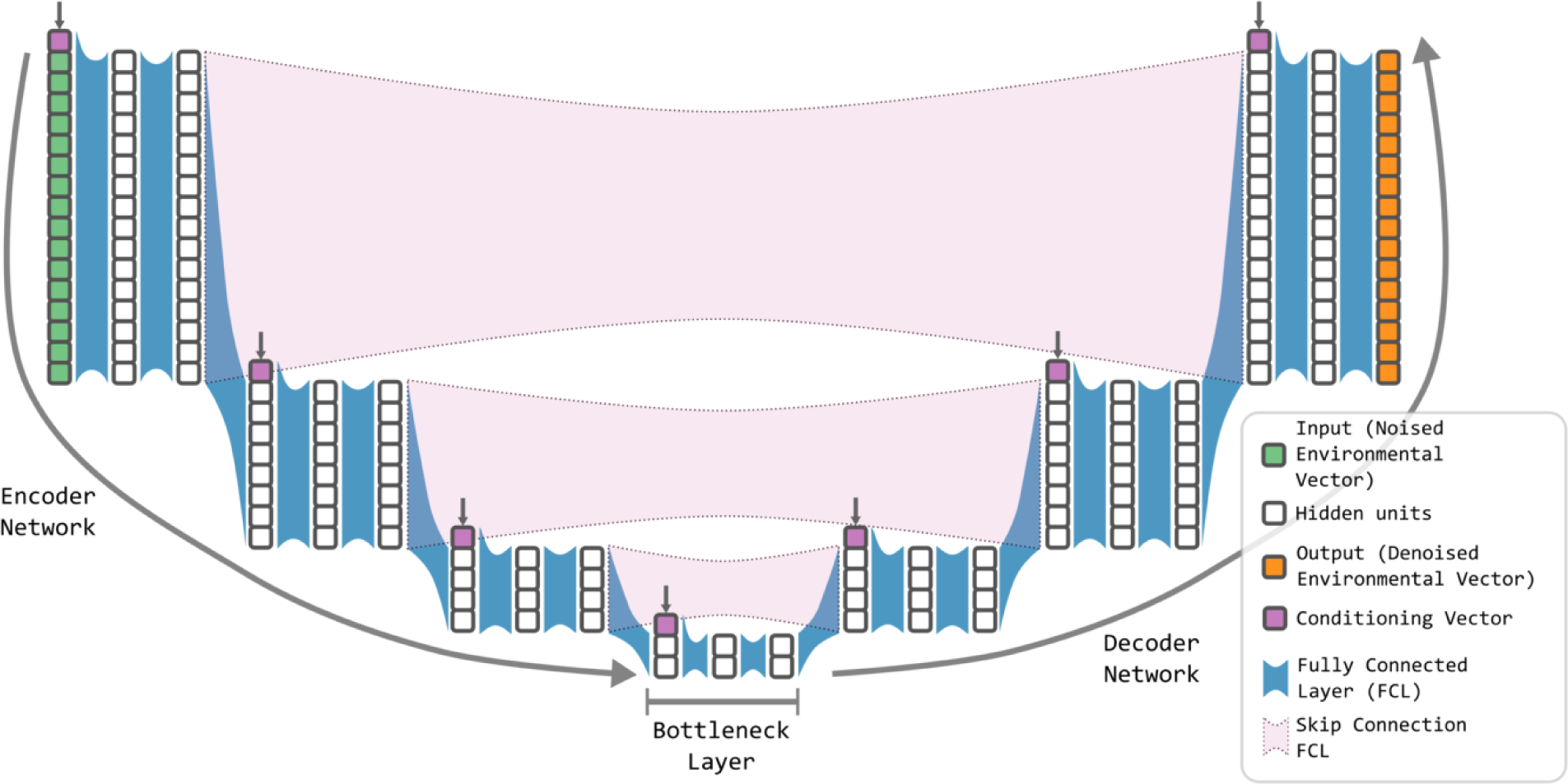
Schematic representation of the modified U-Net architecture used in the Rectified Flow model. The input to the network is a noised environmental vector (green), which undergoes a series of transformations through fully connected layers (blue blocks). The U-Net structure includes downsampling and upsampling paths, with hidden units (gray) processed at each layer. Skip connections (purple dashed lines) preserve feature information between corresponding layers, enhancing the model’s ability to capture multi-scale patterns in the data. Conditioning vectors (pink) provide species-specific context at multiple stages, integrating key environmental and biological factors into the transformation. The output (orange) is the denoised environmental vector, representing a structured transformation from noise to the target distribution, guided by the learned vector field. This architecture supports efficient and accurate sampling within the Rectified Flow model, leveraging hierarchical feature extraction and integration across the network.

**Figure 3:**
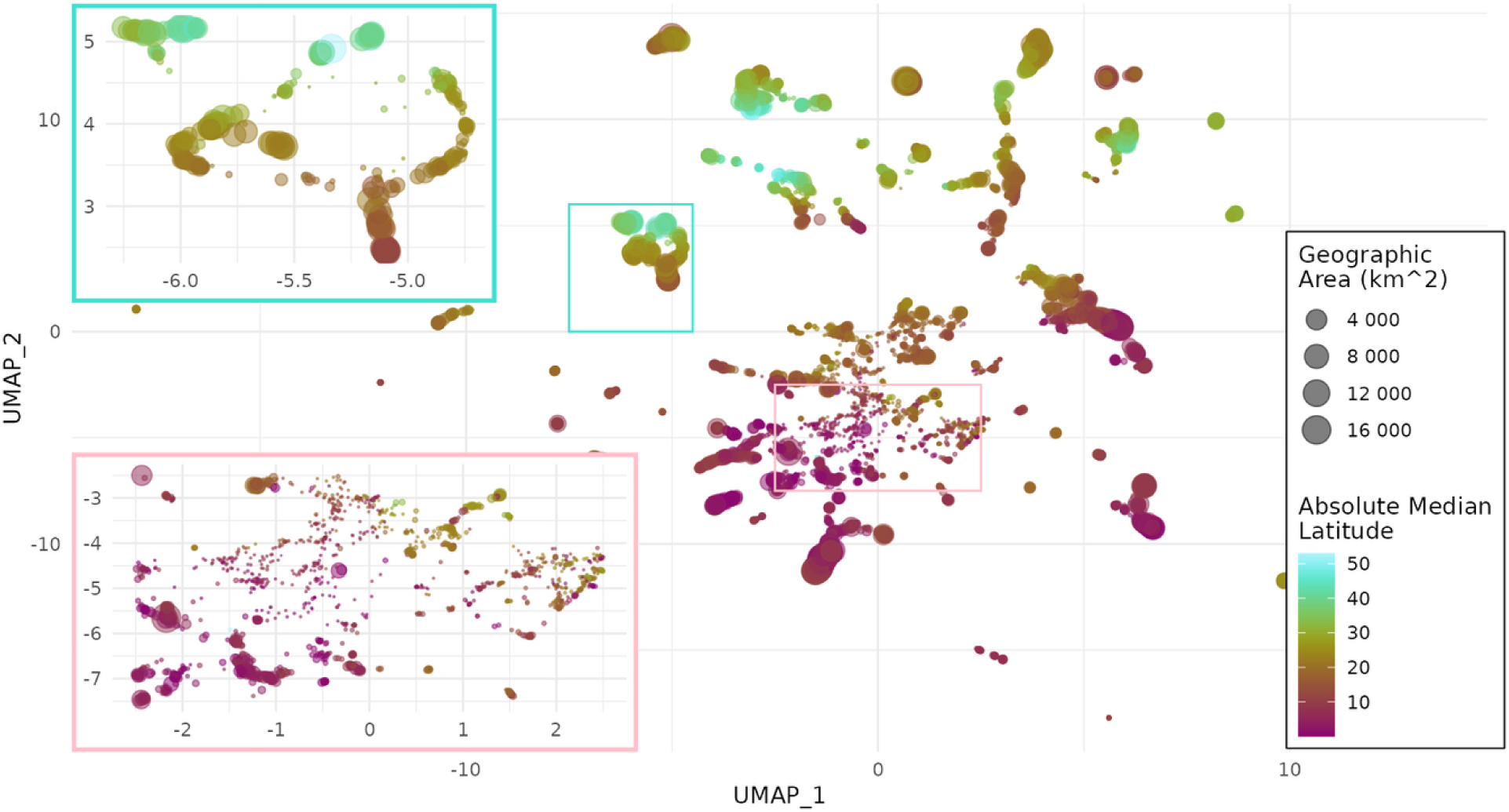
UMAP visualization of the learned latent niche space for reptile species with insets showing zoomed-in regions of interest. Each point represents a species, with its position in the latent space determined by the similarity of its inferred environmental niche. The color gradient indicates the absolute median latitude of each species’ geographic range, with cooler colors representing species closer to the equator and warmer colors representing species at higher latitudes. Point size corresponds to the species’ geographic range area, with larger points indicating larger ranges. The colored rectangles on the main plot correspond to zoomed-in regions displayed as insets to the left, which show greater detail of clustered species within the latent space. These clusters reveal groups of species with similar ecological niches, despite differences in their geographic regions or range sizes. This is a caption.

### Input Conditioning and Encoding

In addition to the U-net structure, GeODE includes specialized encoding layers for the time variable *t* and environmental conditioning vectors. A linear layer encodes the time step, representing the interpolation factor between noise and target coordinates. Another linear layer processes the environmental vectors, embedding them into a latent space that informs the transformation from noise to geographic coordinates. These encoded time and environmental vectors are concatenated with the latent representations from the U-net, allowing the model to incorporate both spatial and environmental dependencies into its predictions.

### Training Data Creation

Training data for GeODE is generated through a Monte Carlo sampling process. Gaussian noise samples are drawn for both the latitude and longitude dimensions, creating initial random coordinate sets. These coordinates are linearly interpolated with target coordinates (actual occurrence points), guided by the ODE. This interpolation path forms the input for training, allowing the model to learn how to evolve from noise to realistic geographic distributions.

### Model Training and Implementation

GeODE was implemented in R using the torch package, leveraging GPU acceleration with CUDA for efficient training.

The full model that combines NichEncoder and GeODE to generate species distribution models was named the **NicheFlow** model. It requires the training of 5 generative models that are chained together. NichEncoder is composed of three models:

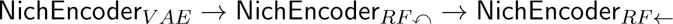

Where *V AE* refers to the initial Variational Autoencoder model, *RF-r--* refers to the stage 1 Rectified Flow model and *RF ←* refers to the stage 2 Rectified Flow model, which has had its ODE rectified (made linear). GeODE is composed of two models:

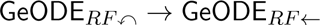

These three models are chained and each needs to be trained on the output of the model to its immediate left. This means that NichEncoder and GeODE can be trained in parallel, but the sub-models have to be trained sequentially. I sequentially trained each of the sub-models in NichEncoder and GeODE (in parallel), each on a Nvidia A100 GPU. Once trained the *RF-r--* models could be discarded and the *RF ←* used for the rectified flow part of the model. These models are much more computationally efficient because they have been ‘rectified’, meaning they can be well approximated by only a single step of ODE integration.

For the *z*_species_ latent space we set the dimension to 32. If fewer dimensions were needed the L2 penalty apply to the loss would shrink some dimensions to effectively zero variance.

Utilizing multiple A100 GPUs to parallelize model training where it was possible, all 5 models that need to be trained to make up *NicheFlow* took less than 1 week to train in total for 2,500 epochs, 6,000 epochs, 3,000 epochs, 5,000 epochs, and 2,000 epochs for NichEncoder*_V AE_*, NichEncoder*_RF-r--_*, NichEncoder*_RF ←_*, GeODE*_RF-r--_*, and GeODE*_RF ←_*, respectively.

### Model Evaluation

To evaluate the performance of the generative species distribution model (SDM), I compared model predictions to observed test occurrence points using hexagonal binning and spatial aggregation. This approach allowed us to transform the model’s generative output into a comparable format for calculating standard SDM performance metrics, such as accuracy, ROC-AUC, and True Skill Statistic (TSS), facilitating comparisons with other SDM approaches.

I tested the performance on 424 randomly selected species from the °10,000 in the training dataset. The random sampling was stratified by data deficiency (fewshot) status, three levels of geographic range size (2.5 - 25 km2, 25 - 220 km2, and 220 - 2000 km2), and three levels of absolute mid-latitude (0 - 17 degrees, 17 - 34 degrees, 34 - 51 degrees), to get a geographically representative set of species. No reptile species had an absolute mid-latitude greater than 51 degrees.

Hexagonal binning was used to approximate the probability of species occurrence across the study area. The geographic predictions of the model, consisting of sampled points, were grouped into hexagonal grid cells, creating an occurrence density map. The relative occurrence probability within each hex cell was calculated as the proportion of predicted points within that hex compared to the total predicted points across all hexes. This procedure allows the generative output, which produces samples rather than explicit probabilities, to be converted into a spatially aggregated form that is comparable to traditional SDMs that produce per-cell probabilities.

The evaluation was conducted within a more localized geographic context, which is a common approach in traditional SDMs that typically model distributions within a “background area” — a region of interest surrounding the species’ known occurrence points. While NicheFlow, being a global model, is trained to localize species distributions across the entire world, I confined its predictions to a smaller geographic region to simulate the background area used in traditional SDMs. Specifically, I identified the set of ecoregions overlapping the species’ known occurrence points and used these ecoregions as the background area for evaluation. This approach allowed us to evaluate whether the model could accurately localize the species within its natural ecoregions, which is a more fine-grained task compared to merely determining the part of the world where the species is likely to occur.

I compared the predicted occurrence probabilities within these localized ecoregions to observed occurrence points. For each hexagonal cell, I calculated the proportion of observed occurrence points (from test data) and used this as the “true” occurrence probability. The True Skill Statistic (TSS), also known as Youden’s J-index (Youden, 1950), was used as the primary evaluation metric to assess the model’s ability to differentiate between presence and absence cells.

In addition to TSS, I calculated several other standard SDM evaluation metrics, including Accuracy, ROC-AUC, and F-measure. Accuracy is the proportion of correctly predicted presences and absences across all hexes. ROC-AUC (Receiver Operating Characteristic - Area Under Curve) quantifies the model’s ability to discriminate between presence and absence, with values closer to 1 indicating better discrimination. F-measure balances precision and recall in binary classification problems, providing a robust metric for evaluating the presence/absence predictions across hexes.

To calibrate the model predictions, I applied a thresholding procedure (Phillips et al., 2006). The generative model outputs continuous probabilities for each hexagonal cell, so a threshold is needed to convert these probabilities into binary presence/absence predictions. I applied a threshold optimization approach based on TSS, selecting the threshold that maximizes the TSS score for the test data. This allowed us to determine the optimal cutoff for classifying a cell as occupied or not, improving model interpretability and comparison with other SDM methods.

The evaluation process was implemented using the tidymodels framework (Kuhn and Wickham, 2020) for calculating metrics and the probably package (Vaughan, 2020) for threshold optimization. Geographic data manipulation and visualization were performed using the sf (Pebesma, 2018) and h3 (Brodrick, 2019) packages, ensuring accurate spatial alignment and efficient processing of hexagonal grids.

### Dataset

#### Species Distribution Data

The dataset used to test the model consists of species distribution maps for 10,064 extant reptile species, encompassing a wide variety of taxa, including lizards, snakes, turtles, amphisbaenians, and crocodiles (Roll et al., 2017). These species distribution maps represent polygons of the species’ extents of occurrence, which were derived from a combination of sources, including field guides, museum databases, the Global Biodiversity Information Facility (GBIF), the International Union for Conservation of Nature (IUCN), and expert observations. This rich dataset provides comprehensive global coverage of reptile distributions and is well-suited to train generative models for species distribution prediction.

For the purposes of this study, I transformed the polygonal data into point occurrences to better suit the requirements of the generative modeling approach. Using the R package sf (Pebesma, 2018), I uniformly sampled 800 points within each polygon to serve as the main training dataset. Additionally, I created a held-out test set for each species by sampling a further 400 points, which were excluded during training and used to evaluate the model’s performance.

In addition to testing the model on species with abundant occurrence data, I specifically designed a set of species to simulate real-world scenarios where distribution data is sparse. This subset, referred to as the ‘few-shot species,’ includes species for which I only sampled 4 random points from their distribution. This design choice allowed me to evaluate the model’s capacity to learn distributions of species with highly limited data—a situation that is frequently encountered in real biodiversity datasets. The few-shot testing is an important component of evaluating the model’s robustness to data deficiency.

Moreover, a subset of species was deliberately left out of the training set entirely to test the model’s zero-shot capabilities, as described in previous sections. This experimental design allows for a comprehensive evaluation of the model’s ability to predict species distributions across a wide spectrum of data availability, from well-sampled species to those for which no prior occurrence data was used during training.

### Environmental Data

In this study, I utilized the CHELSA-BIOCLIM dataset (Karger et al. 2017) to extract 32 environmental variables crucial for species distribution modeling. These bioclimatic variables, which include mean annual temperature, precipitation patterns, and seasonality, provide insights into the climatic factors that shape species distributions (Karger et al., 2017). The high spatial resolution of 30 arc-seconds (°1 km²) in the CHELSA-BIOCLIM dataset enables precise mapping of environmental conditions at species’ occurrence points, which is particularly useful in ecological niche modeling. Due to large amounts of missing data in 2 of the 32 CHELSA-BIOCLIM variables (), these were subsequently dropped from the training data used by NicheFlow.

To integrate the environmental data into the model, I used the terra package in R (Hijmans, 2022) to extract these variables at specific spatial points corresponding to species occurrence locations.

## Results

### NicheFlow captures a representation of niches

After model training, I found 2 of the 32 dimensions that the model were initialized with shrank to near zero variance during training so the effective dimension of the resulting latent species niche space was 30. To visualize the structure of this latent niche space I used the UMAP algorithm (McInnes et al., 2018). UMAP (Uniform Manifold Approximation and Projection) is a dimensionality reduction technique that helps visualize complex, high-dimensional data in two or three dimensions, while preserving important structure and relationships between data points. It is widely used in biology for tasks such as visualizing gene expression patterns, clustering species based on traits, or analyzing ecological datasets. UMAP is particularly valued for its ability to capture both local and global data patterns more effectively than older methods like PCA or t-SNE. I used it to reduce the 30 effective dimensions of the niche space to 2 for easy visualization (Figure \ref*{*985500*}*)

I found that species were in some case widely separated in the two UMAP axes, appearing in multiple clusters throughout the space. I also found some association between the UMAP space and the total range size of the species being modeled as well as it median latitude (Figure \ref*{*985500*}*). More specifically I found that latitude separated species in the UMAP space, in this case with high latitude species tending to be at high values of the second UMAP axis, whereas low latitude species tended to have low values of UMAP 2. On the other hand, species with small ranges tended to be toward the middle of the UMAP space, and larger ranged species towards the edges, forming a halo around the smaller ranged species. This suggests the latent niche space has captured something ecologically meaningful in it’s vectors. Further exploration of the meaning of these niche vectors will be conducted in a follow-up study.

### Model evaluation metrics show NicheFlow captures geographic distributions accurately

The performance of the NicheFlow model was evaluated across two key scenarios: species with abundant data and ‘few-shot’ species, where only four occurrence points were used for training. The AUC metric served as the primary evaluation metric, with F-score and TSS results displaying similar trends. All evaluation metrics were calculated based on a held-out sample of 400 test points per species, including for the few-shot species. This consistent test sample size allowed for a robust comparison between different data abundance scenarios.

For data-abundant species, the model exhibited strong predictive accuracy, particularly for species with small and medium geographic ranges (Figure \ref*{*751151*}*). Examples of environmental and geographic predictions for a randomly chosen data-abundant species can be seen in Figures 5 and 6. For evaluation metrics, at high latitudes, small-range species achieved the highest mean AUC (0.99 ± 0.01). However, performance for large-range species was lower across all latitudinal zones, with a notable dip at equatorial latitudes (0.75 ± 0.02).

**Figure 4:**
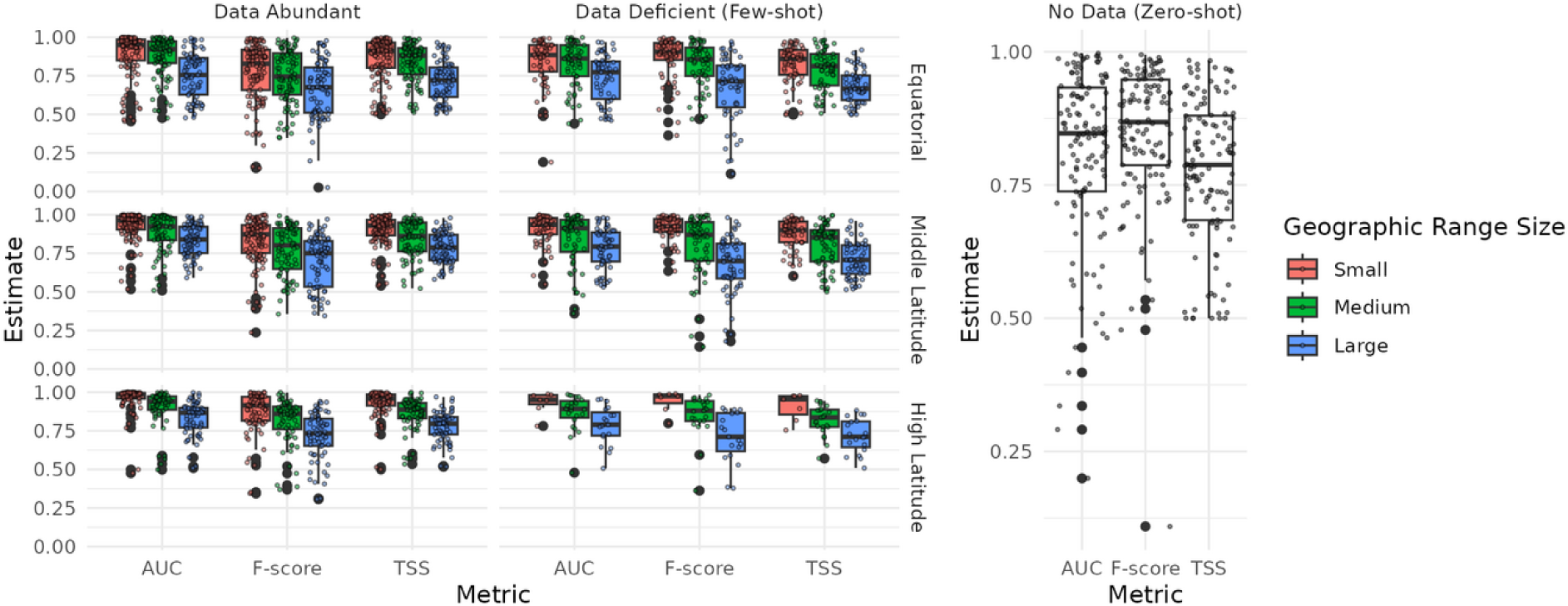
Evaluation of NicheFlow model performance across species with different geographic range sizes and data availability levels, measured using AUC, F-score, and TSS metrics. The left-hand panels depict results for species with abundant occurrence data, while the right-hand panels focus on ‘few-shot’ species, for which only 4 training points were provided. Results are further stratified by latitudinal zone (Equatorial, Middle Latitude, High Latitude) and geographic range size (Small, Medium, Large). Each boxplot summarizes the distribution of the given metric across species, with higher values indicating better performance. Note that TSS has been normalized to fall between 0 and 1 to facilitate comparison with the other metrics (normally it ranges between −1 and 1).

**Figure 5:**
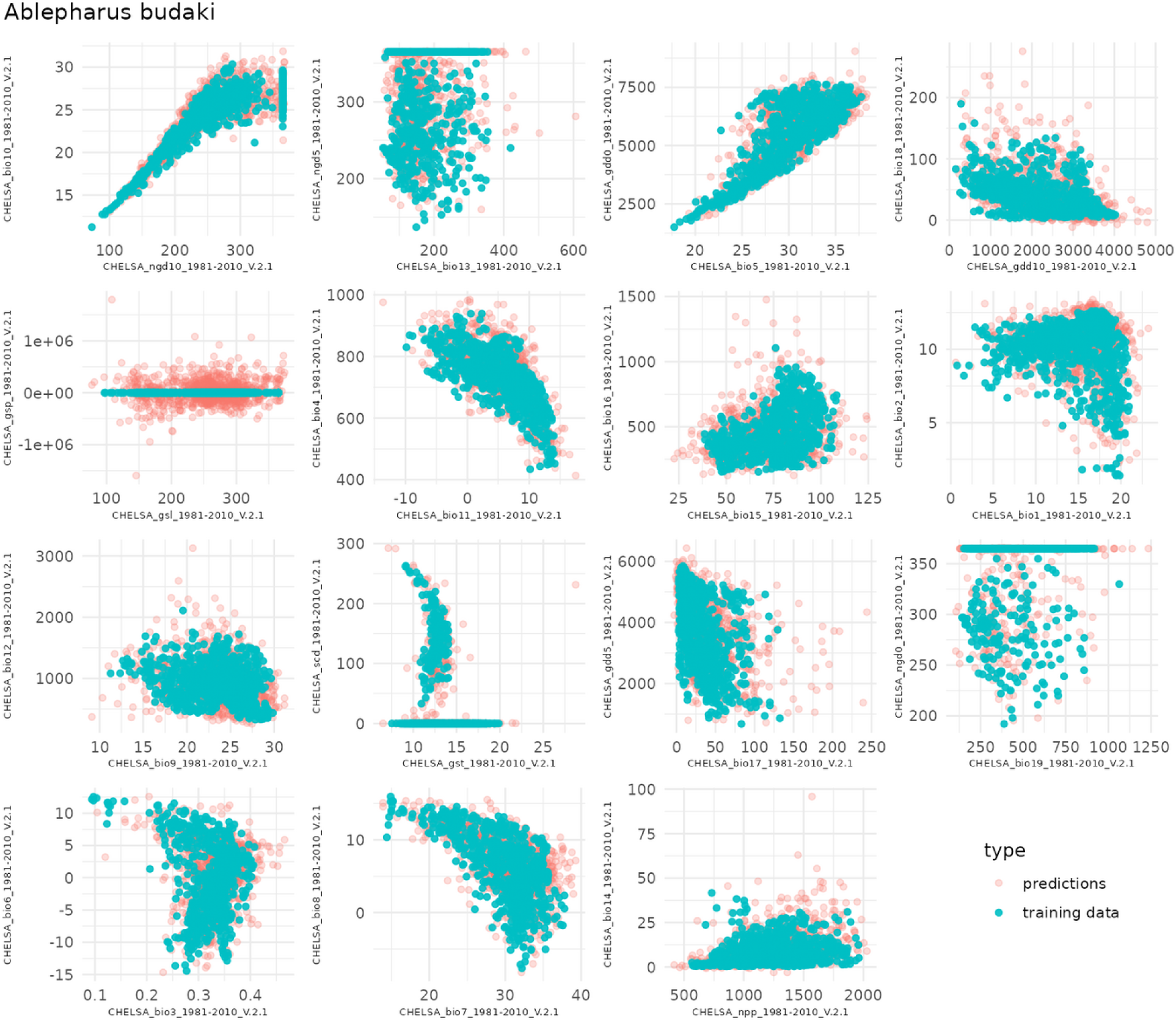
Environmental niche predictions for *Ablepharus budaki* showing comparisons between predicted and training data across 15 pairs of bioclimatic variables from the CHELSA dataset. Each scatterplot compares the environmental variable’s predicted values (red) to the training data (blue). The model shows good alignment between predicted and observed environmental variables, demonstrating how well the model captures the environmental space associated with the species.

**Figure 6:**
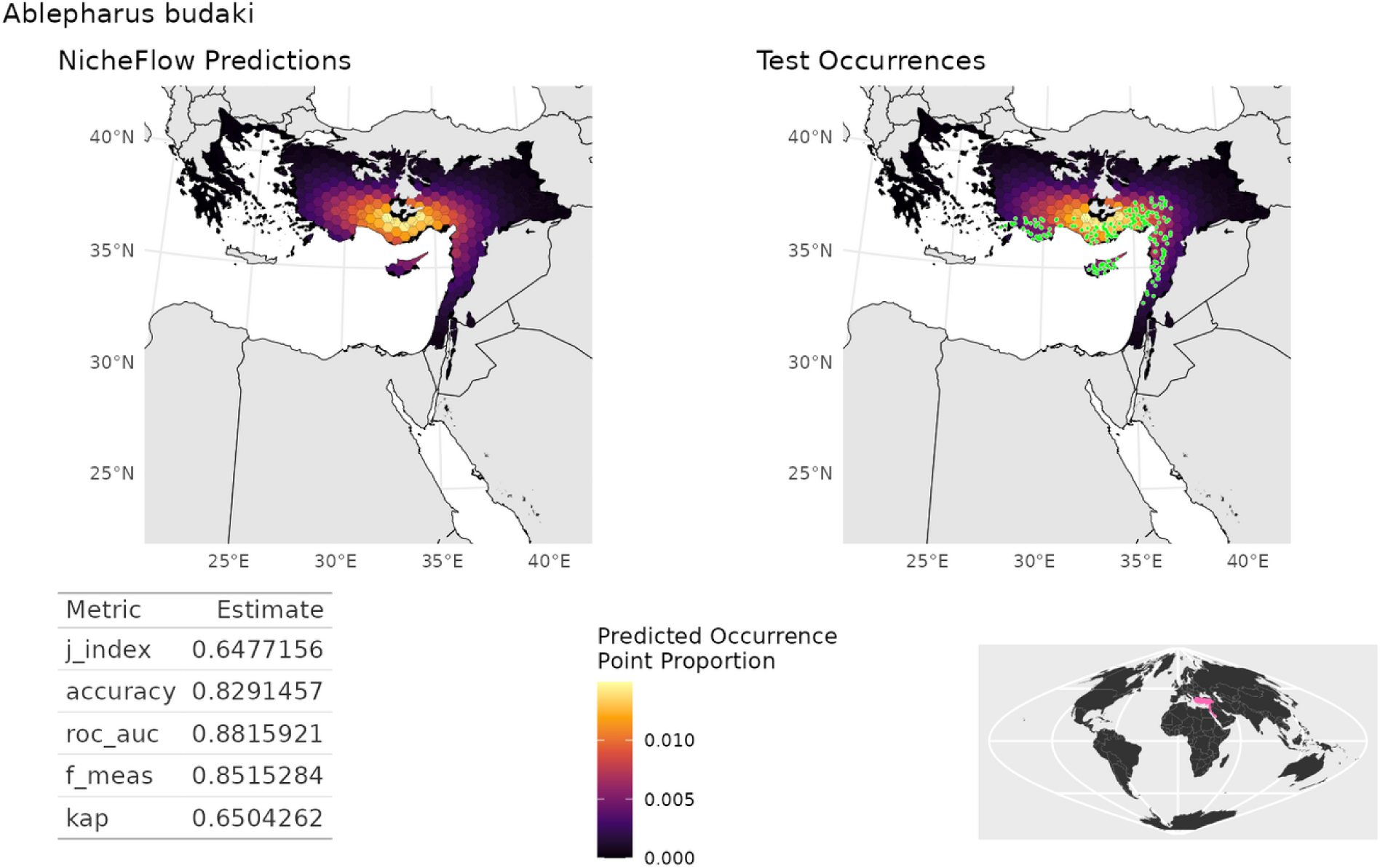
Geographic prediction maps for *Ablepharus budaki* comparing NicheFlow predictions to the species’ true test occurrences. The left panel shows the predicted occurrence probability in hexagon bins across the species’ range, while the right panel depicts the test occurrence points used for evaluation. The table below the maps summarizes the model performance metrics, with an AUC of 0.88 indicating strong predictive accuracy for this species’ distribution. The inset globe highlights the species’ location within its global context.

In the few-shot species scenario, where the model was trained on only four occurrence points, its performance remained impressive. Examples of environmental and geographic predictions for a randomly chosen data-abundant species can be seen in Figures 7 and 8. AUC for small-range species at high latitudes achieved a value of 0.95 ± 0.01. AUC values were also particularly high for small and medium-range species in middle latitudes (0.94 ± 0.01 and 0.91 ± 0.02, respectively). However, as seen in the data-abundant species, large-range species at equatorial latitudes exhibited the lowest AUC performance (0.77 ± 0.03). The consistently strong performance, even with few-shot training data, demonstrates the robustness of NicheFlow in making accurate predictions for under-sampled species.

**Figure 7:**
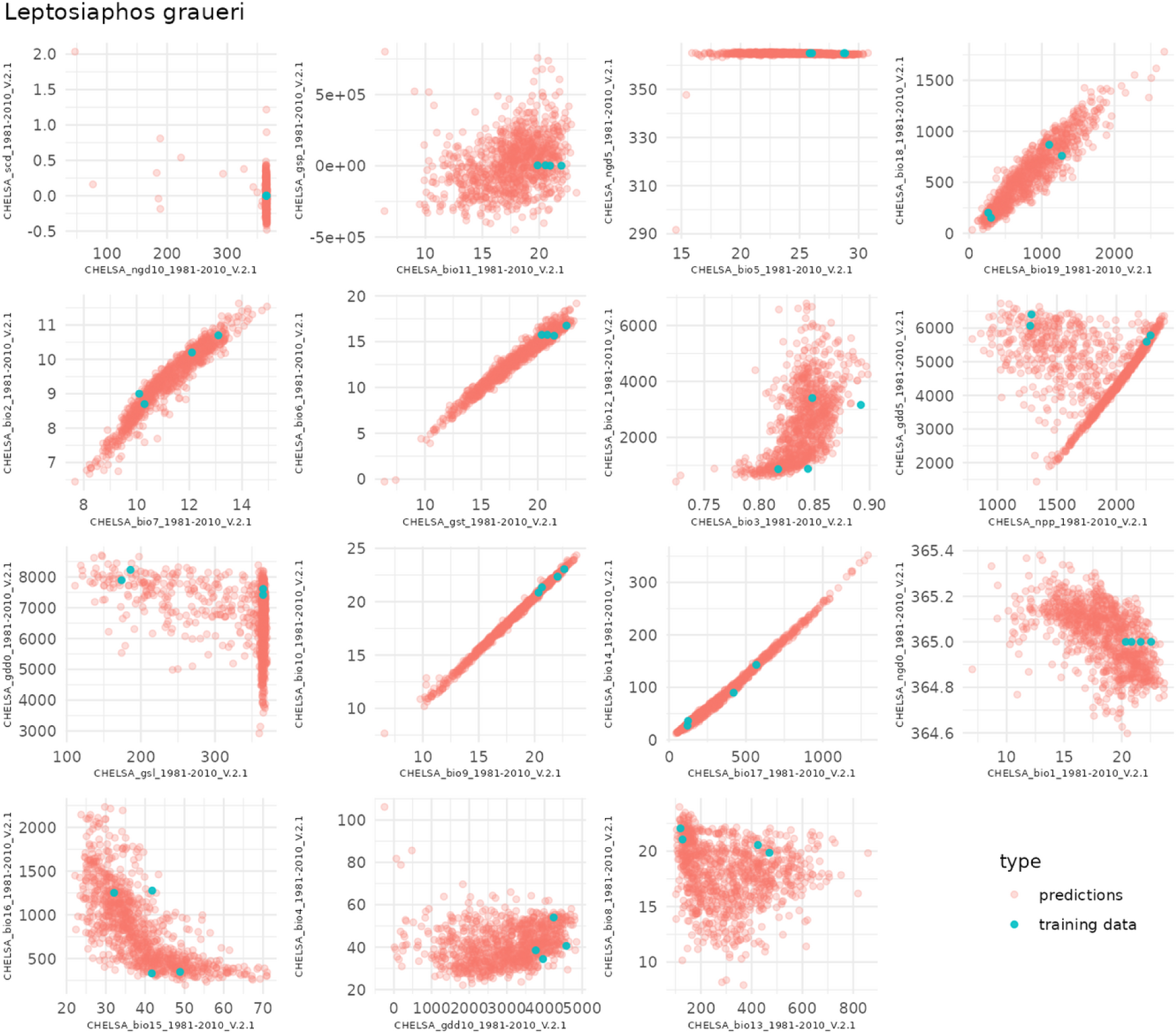
Plots comparing model predictions and observed occurrences in environmental space for the species *Leptosiaphos graueri*, a few-shot species with only 4 training points. Pairwise scatterplots comparing the predicted environmental variables (red) to the true occurrence data (blue). Each panel represents a different combination of 16 environmental variables sampled from the CHELSA-BIOCLIM dataset, allowing for the evaluation of the model’s ability to replicate the environmental conditions associated with the species’ range. This plot highlights the model’s performance, particularly for species with extremely limited training data.

**Figure 8:**
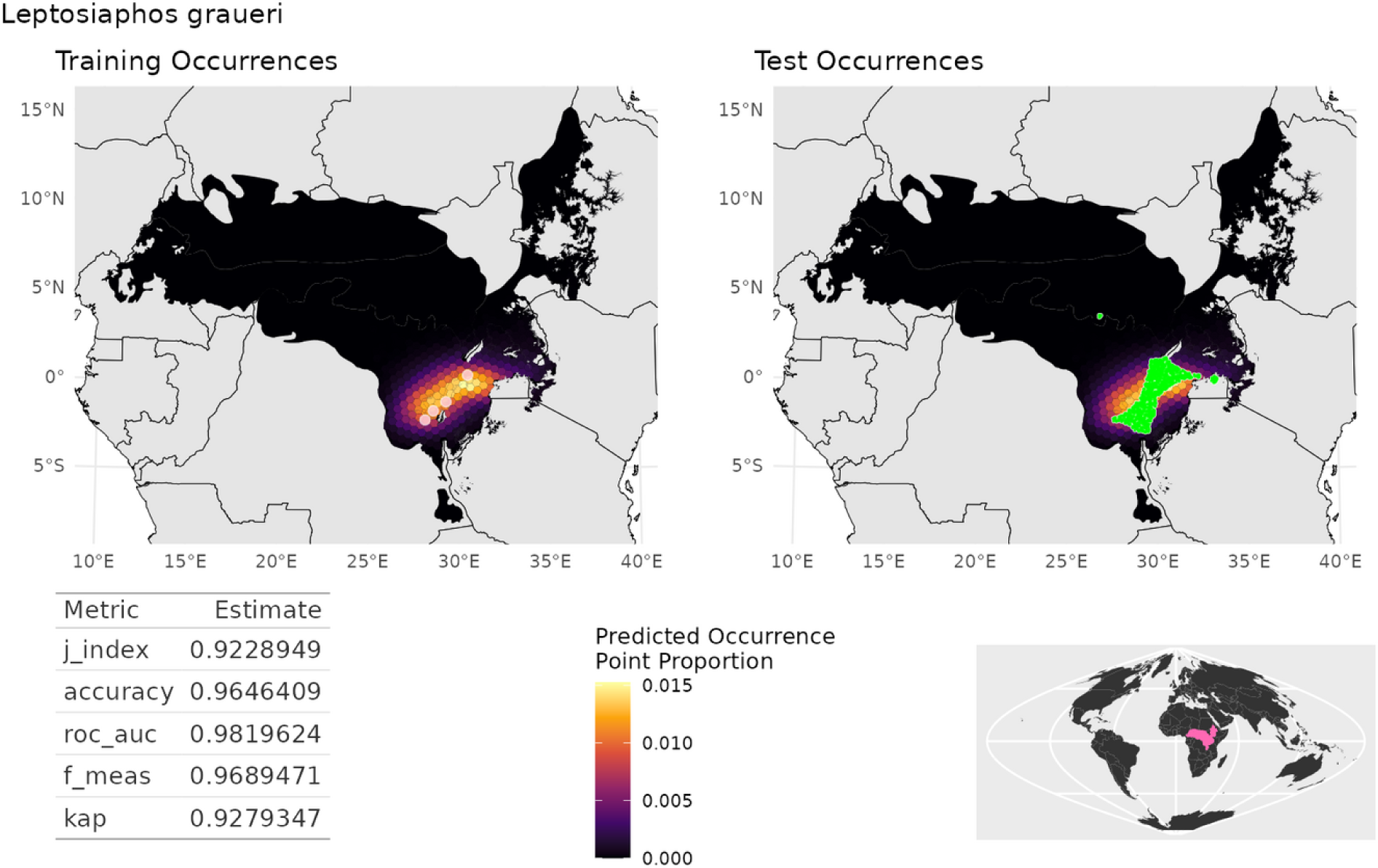
Maps comparing model predictions and test occurrences for the species Leptosiaphos graueri, a few-shot species with only 4 training points. The left panel shows the hex-binned predictions from the NicheFlow model along with the four training points in yellow, while the right panel shows the actual test occurrences (400 points). Colors indicate predicted occurrence proportions for each hexagon. The table below provides key evaluation metrics, including J-index, accuracy, ROC-AUC, F-measure, and True Skill Statistic (J-index). The inset map shows the global context for the region where this species occurs. Predictions somewhat underestimate the true extent of the range, a common occurrence for data-deficient species and probably a result of the few randomly sample location being more likely to come from the centre of the range. Nevertheless evaluation metric are very good with AUC of 0.98.

The lower performance observed for large-range species is likely attributable to the generative sampling strategy. Large-range species require more points to adequately capture the full extent of their distribution. With the current fixed sampling approach, some hexagonal grid cells that encompass the large-range species may contain zero points due to random chance. This results in sparse geographic coverage, limiting the accuracy of predictions for large-range species. In future work, I plan to address this issue by adaptively sampling more points for large-range species, iteratively sampling until cell frequencies converge to a stable value. This will ensure more comprehensive coverage of large ranges, especially at equatorial latitudes, where environmental heterogeneity demands more extensive sampling to accurately represent species distributions. This strategy is expected to improve the model’s accuracy for species with expansive distributions.

Across all species, the model showed robust performance even for few-shot species, where only four training points were available, compared with 800 points for all other species. Specifically, the average AUC for data-deficient species was 0.87, while data-abundant species achieved a slightly higher average AUC of 0.92. Interestingly, few-shot species exhibited a higher F-score of 0.86 compared to 0.81 for data-abundant species, suggesting that the model effectively captured the general characteristics of the species distributions despite extreme data deficiency. The TSS values for few-shot species, although lower, still indicate a reasonable ability to differentiate presence from absence in the test data. This demonstrates the model’s capability of learning useful species-environment relationships, even in highly data-scarce situations.

This held-out test set consisted of 400 points for both data-abundant and few-shot species, providing a reliable evaluation of the model’s predictive capacity across different data regimes. The model’s generalization ability, particularly for species with very limited occurrence records, underscores its potential for addressing real-world biodiversity data challenges, where species are often data-deficient.

### NicheFlow successfully performs Zero-Shot prediction

Even with species that had no data in the training set, it was possible to get good quality distribution prediction from NicheFlow by match occurrence point of the species to generated occurrence point distribution from the model and using this to optimize the zero-shot species latent niche space vector z species (Figure 9). Overall I tested 124 species that had been held-out entirely from the training set (Figure 4, right panel). When tested against the 400 held-out occurrence points, on average NicheFlow predicted species distribution had an AUC of 0.81 ± 0.01 (median = 0.84). This is substantially lower than for data abundant or few-shot species but nevertheless remarkable considering the training sample size of N = 0. There was also more spread for zero-shot species, with them being the only species to occasionally exhibit an AUC less than 0.5, representing predictions that were worse than random. This most likely occurred as a result of the latent vector optimization failing to find a good optimum, either because a good optimum did not exist in the latent space, or more likely because it got stuck in a local optimum in a rough loss landscape.

**Figure 9:**
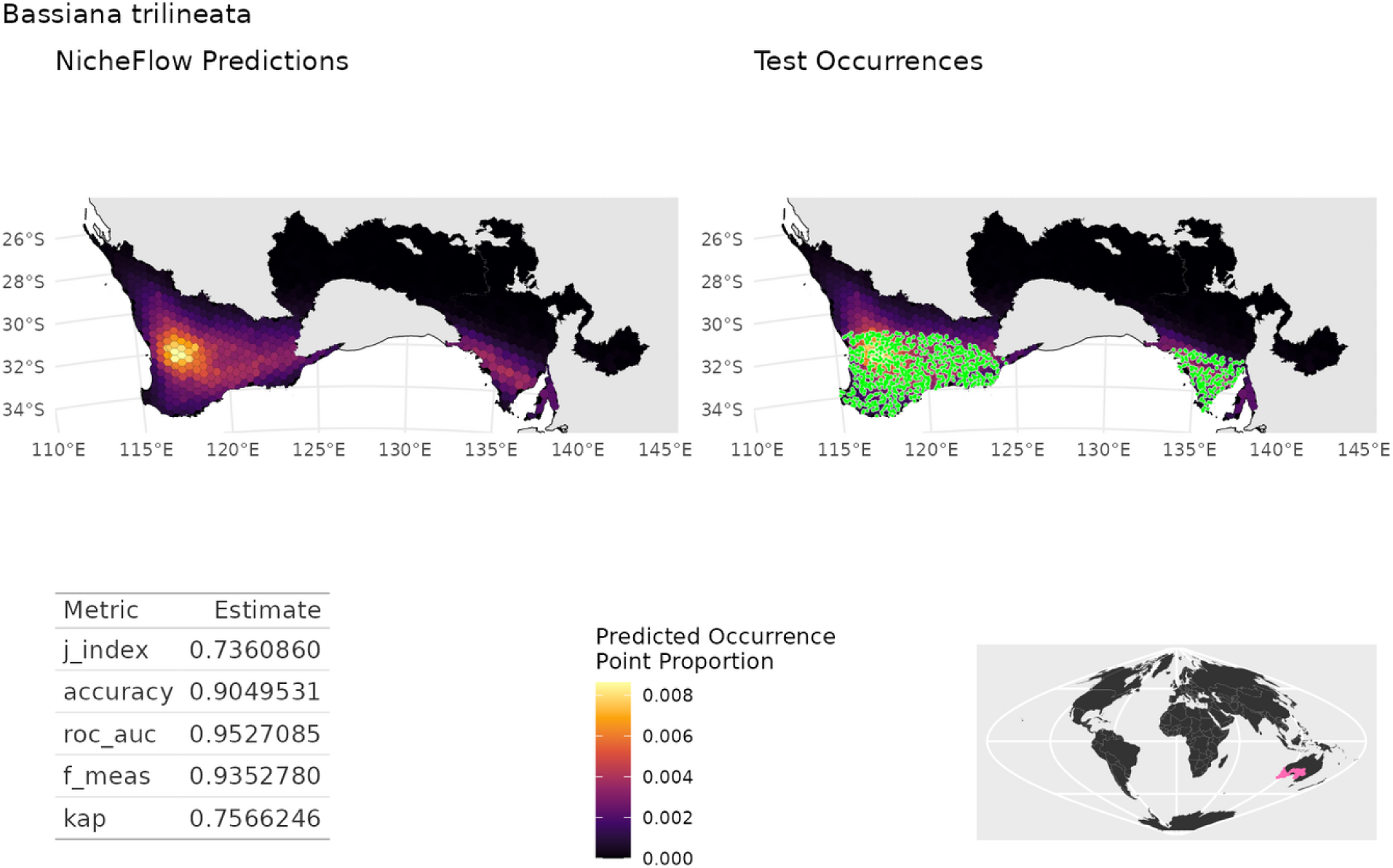
Zero-shot geographic predictions for Bassiana trilineata. The NicheFlow model’s predicted occurrence density is shown on the left, derived entirely through zero-shot learning without training data for this species. Hexagonal bins represent the proportion of predicted occurrences, with brighter hexes indicating areas of higher predicted density. Test occurrences, shown on the right in green, are overlaid for comparison to the model’s predicted points. The species’ range is accurately captured despite the absence of direct training data, as reflected in high evaluation metrics, including an AUC of 0.95, F-measure of 0.93, and a True Skill Statistic (J-index) of 0.74. The bottom right inset shows the species’ geographic location.

## Discussion

### Advancing Species Distribution Modeling with Foundation Models

NicheFlow represents a significant leap forward in species distribution modeling (SDM), harnessing the power of generative AI to tackle long-standing challenges in ecological predictions. By employing a flexible architecture capable of generalizing across species and ecosystems, NicheFlow has the potential to revolutionize how we model, understand, and conserve biodiversity -- a potential foundation model for ecology (Bommasani et al., 2021).

The application of foundation models in ecology couldn’t be more timely. Traditional SDMs have long grappled with limited and biased data, particularly the absence of true absence data (Elith et al., 2006). NicheFlow addresses this challenge head-on by integrating species embeddings, allowing for “strength sharing” between species. This innovative approach enhances predictions for rare or data-limited species, building upon joint species distribution models (JSDMs) that leverage species correlations (Warton et al., 2015; Pollock et al., 2014; Ovaskainen et al., 2016). However, NicheFlow goes a step further, enabling non-linear generalization and thus capturing more complex ecological relationships.

One of NicheFlow’s key strengths lies in its ability to extract patterns from large, heterogeneous datasets. This capability could provide a transferable understanding of niche space, enabling predictions in new regions or under future climate scenarios. Once trained and released the power of the model can be utilized or fine-tuned by anyone in the research or practitioner community. Such transferability and share-ability aligns perfectly with growing calls for open science and data sharing in ecology (McKiernan et al., 2016; Hampton et al., 2015), extending it beyond data to model too, and paving the way for more collaborative and comprehensive computational ecology research.

### Generative Approach: A Paradigm Shift in SDM

NicheFlow marks a paradigm shift in species distribution modeling. Unlike traditional SDMs that operate within a discriminative framework (Guisan & Thuiller, 2005; Franklin, 2010; Araújo & Peterson, 2012), NicheFlow explicitly models the conditional distribution of species in environmental space, an approach with some similarities to environmental density estimation methods like hypervolume (Blonder et al. 2018), but using a generative multi-species approach (see Supporting Information for a detailed discussion of connections between NicheFlow and other SDM approaches). The generative approach of NicheFlow offers significant advantages, particularly in handling novel climates and predicting species responses to changing conditions (Araújo & Rahbek, 2006; Warren et al., 2014).

Perhaps the most remarkable outcome of this approach is NicheFlow’s effectiveness in predicting distributions for data-deficient or few-shot species. Few-shot learning, the ability to generalize with limited examples (Wang et al., 2020), is crucial in ecology where many species have sparse occurrence records (Breiner et al., 2015). NicheFlow’s latent niche space allows it to leverage patterns learned from data-rich species to benefit data-deficient ones. The result? Robust predictions (average AUC *>* 0.85) for few-shot species, a feat that traditional SDMs often struggle to achieve.

Taking this a step further, NicheFlow demonstrates potential for zero-shot learning, predicting distributions for species entirely absent from the training data. This capability extends the model’s utility dramatically, allowing researchers and practitioners to use it without retraining, regardless of data availability.

### Addressing Climate Change and Conservation Challenges

In the face of rapid climate change, NicheFlow’s flexibility in simulating species responses under novel conditions offers a significant advantage. Traditional SDMs often struggle with non-analog climates (Williams & Jackson, 2007), but NicheFlow’s generative approach may better capture species’ potential responses to new environmental combinations. This capability could prove invaluable in identifying future suitable habitats for species reintroductions or in conservation planning (Guisan et al., 2013; Hannah et al., 2007).

Moreover, NicheFlow’s joint species distribution capabilities provide a powerful tool for community-level conservation planning. By modeling multiple species simultaneously, we can identify high-biodiversity regions or at-risk species assemblages more effectively (Pereira et al., 2010). This aligns perfectly with global biodiversity initiatives aiming to preserve ecosystem integrity (Convention on Biological Diversity, 2021), offering a more holistic approach to conservation.

### New Frontiers in Niche Theory and Community Ecology

NicheFlow’s architecture opens up exciting new avenues for exploring fundamental questions in niche theory. Its ability to capture complex, non-linear relationships in high-dimensional environmental space aligns beautifully with Hutchinson’s n-dimensional hypervolume concept (Hutchinson, 1957; Blonder, 2018; Holt, 2009). By examining the learned embedding space, we could gain unprecedented insights into niche dimensionality, breadth, overlap, and evolution across taxa.

The model’s capacity to generate samples from species’ environmental niches enables novel approaches to studying niche dynamics. This could reveal patterns of niche conservatism or divergence (Wiens et al., 2010; Pearman et al., 2008), shedding light on long-standing questions in evolutionary ecology. Furthermore, it could facilitate exploration of community assembly processes, allowing us to test hypotheses about environmental filtering versus competitive exclusion (Kraft et al., 2015; Cadotte & Tucker, 2017) with greater precision than ever before.

NicheFlow’s ability to generate hypothetical species distributions based on interpolations in the embedding space opens up fascinating possibilities for evolutionary research. We could simulate potential distributions of hybrid species or explore “empty niche space”(Schluter, 2000), providing new insights into adaptive radiation and niche evolution. By combining NicheFlow with ancestral niche reconstruction techniques, we could even predict historical species distributions, offering new avenues for testing biogeographic and niche evolution hypotheses (Wiens & Graham, 2005; Crisp & Cook, 2012; Kozak & Wiens, 2006).

### Caveats and Future Directions

Despite its advancements, NicheFlow is not without limitations. The quality and biases of input data, whether from expert range maps or occurrence records, can significantly impact model outcomes (Hurlbert & Jetz, 2007; Newbold, 2010; Hijmans et al., 2000; Reddy & Davalos, 2003). To address this, future iterations of NicheFlow should leverage multiple data types, creating more comprehensive and nuanced representations of species distributions.

Interpretability remains a challenge, as with many deep learning models in ecology (Merow et al., 2014; Olden et al., 2008). To enhance NicheFlow’s utility for ecological insight, we must focus on improving model interpretability. This could involve incorporating explainable AI techniques or developing methods to translate learned embeddings into ecologically meaningful concepts.

To provide a more nuanced view of species’ ecological niches, it will be critical to better incorporate uncertainty into NicheFlow. We can achieve this by implementing a variational autoencoder variant to model the latent space, facilitating better uncertainty quantification in model predictions. This probabilistic treatment will also enable more effective amortized inference, potentially improving computational efficiency.

Enhancing zero-shot prediction capabilities represents another key area for improvement. By increasing latent space regularization and incorporating auxiliary predictors such as phylogenetic information, species traits, and environmental data, we can significantly expand NicheFlow’s utility in predicting distributions for rare, newly discovered, or data-deficient species.

To truly realize the potential of a foundation model in ecology, we aim to train NicheFlow on distribution data for all terrestrial vertebrates in the next phase of development. This comprehensive dataset will allow the model to capture a wider range of ecological niches and biogeographic patterns, enabling more robust exploration of macroecological patterns and cross-taxa comparisons.

### Ethical Considerations

As we advance this powerful tool, we must not overlook important ethical and societal considerations. Issues of data privacy and ownership, particularly for data from indigenous communities or citizen scientists, necessitate clear guidelines on data usage and sharing (Groom et al., 2017). We must also carefully consider how to share and use model outputs to prevent potential misuse, such as exploitation by poachers or land grabbers.

Ensuring equitable access to NicheFlow is crucial. We must address potential exacerbation of existing inequalities in ecological research and conservation planning due to computational resource requirements. By democratizing access to this advanced tool, we can foster more inclusive and comprehensive global biodiversity research and conservation efforts.

## Conclusion

NicheFlow represents a significant step forward in species distribution modeling, offering new insights into ecological niches and species distributions. As we continue to refine and expand the model, its potential applications in climate change impact assessment, conservation planning, and evolutionary studies are vast. The integration of NicheFlow with other data sources promises to further enhance our understanding of biodiversity patterns and processes, providing crucial tools for addressing mounting ecological challenges in the face of global change. By leveraging the power of foundation models and generative AI, NicheFlow paves the way for a new era in ecological modeling and conservation planning.

## Data Availability

All code used to implement the NicheFlow model is publicly available on Github. Data use to train a proof-of-principle model is availably publicly at https://datadryad.org/stash/dataset/ doi:10.5061/dryad.83s7k and https://chelsa-climate.org/bioclim/

Code for implementing the models is publicly available on GitHub (https://github.com/rdinnager/genAISDM)

## Supporting Information

A supporting information document can be found at https://www.authorea.com/users/5518/articles/1231655-nicheflow-towards-a-foundation-model-for-species-distribution-modelling-supporting-information

This includes an animated figure demonstrating latent niche interpolation for the NicheFlow reptile model.

