## Supporting Information for "NicheFlow: Towards a foundation model for Species Distribution Modelling"

#### Mathematical Details

##### Derivation of the Model Equations

The goal is to estimate the probability distribution of species across geographic coordinates $(X, Y)$, given that the species is $S = s$, and that it occurs ($O_s = 1$) e.g. $P(X, Y|S = s, O_s = 1)$. For simplicity we will use the expression $S = s$ to represent $S = s, O_s = 1$. To include the environment in this probability, this can be represented mathematically as:

$$P(X, Y \mid S = s) = \int_{-\infty}^{\infty} \cdots \int_{-\infty}^{\infty} P(X, Y, \mathbf{E} \mid S = s) \, \mathrm{d}e_1 \cdots \mathrm{d}e_n.$$

This expression introduces the environmental variables $\mathbf{E}$ and integrates over all possible environmental conditions.

1. **Applying the Law of Total Probability**:

   We expand the joint probability using the Law of Total Probability:

   $$P(X, Y \mid S = s) = \int_{-\infty}^{\infty} \cdots \int_{-\infty}^{\infty} P(X, Y \mid S = s, \mathbf{E}) P(\mathbf{E} \mid S = s) \, \mathrm{d}e_1 \cdots \mathrm{d}e_n.$$

   This step decomposes the probability into two components: one that describes how environmental conditions affect the geographic distribution, and another that captures the species' niche, or how environmental conditions influence species occurrence.

2. **Assumption of Conditional Independence**:

   At this point, we make a key biological assumption: the occurrence of a species is driven entirely by environmental conditions, not by geographic coordinates themselves. In statistical terms, this means we assume that species $S$ is conditionally independent of the coordinates $(X, Y)$ given the environmental conditions $\mathbf{E}$. Mathematically, this is expressed as:

   $$P(X, Y \mid S = s, \mathbf{E}) = P(X, Y \mid \mathbf{E}).$$

   This substitution reflects the idea that any effects of geographic coordinates on species occurrence are only through their relationship with the environment. Biologically, this means that the environment determines where species can occur, and coordinates influence species distributions only indirectly via their environmental characteristics. This is related to the standard statistical assumption of "no unmeasured confounders," implying that environmental variables capture all relevant factors affecting species occurrences.

Substituting this assumption, we get:

$$P(X, Y \mid S = s) = \int_{-\infty}^{\infty} \cdots \int_{-\infty}^{\infty} P(X, Y \mid \mathbf{E}) P(\mathbf{E} \mid S = s) \, \mathrm{d}e_1 \cdots \mathrm{d}e_n. \tag{1}$$

3. **Environmental Niche as a Conditional Density with Species Embeddings**:

   The term $P(\mathbf{E} \mid S = s)$ captures the environmental niche of the species, representing a high-dimensional probability distribution of environmental conditions where the species is likely to occur—often referred to as a "hypervolume" in ecological terms. We further decompose this distribution by introducing a latent variable $\mathbf{Z}_s$, which serves as a lower-dimensional vector representation of the species' complex niche. This process is a form of representational learning, where $\mathbf{Z}_s$ is optimized to encapsulate the essential ecological characteristics of the species.

   Specifically:

$$P(\mathbf{E} \mid S = s) = \int_Z P(\mathbf{E} \mid \mathbf{Z} = \mathbf{z}_s) P(\mathbf{Z} = \mathbf{z}_s) \, \mathrm{d}z,$$

   where $\mathbf{Z}_s$ represents the species' niche in a lower-dimensional space, allowing for a more efficient and flexible representation of complex environmental dependencies.

4. **Combining the Integrals**:

   Substituting this into Equation (1), we arrive at the final expression:

$$P(X, Y \mid S = s) = \int_{-\infty}^{\infty} \cdots \int_{-\infty}^{\infty} P(X, Y \mid \mathbf{E}) \left( \int_Z P(\mathbf{E} \mid \mathbf{Z} = \mathbf{z}_s) P(\mathbf{Z}) \, \mathrm{d}z \right) \mathrm{d}e_1 \cdots \mathrm{d}e_n. \tag{2}$$

This final equation (2) is a combination of two probability distributions that are independent of one another:

1. $P(X, Y \mid \mathbf{E})$: Represents the probability of geographic coordinates given environmental conditions.
2. $P(\mathbf{E} \mid S = s)$: Represents the environmental niche of the species, describing how likely a species is to occur under different environmental conditions.

Biologically, $P(\mathbf{E} \mid S = s)$ captures the species' niche, detailing the range of environmental conditions under which a species can thrive. Meanwhile, $P(X, Y \mid \mathbf{E})$ maps these environmental conditions to geographic locations, indicating where such suitable conditions are found on Earth.

#### Distance Metrics for Comparing High Dimensional Distributions

Let $\mathbf{E}_{\text{pred}}$ and $\mathbf{E}_{\text{true}}$ be two matrices where rows represent individual environmental vectors associated with predicted and observed occurrences, respectively. These matrices can have different numbers of rows, reflecting the flexibility of the distance measures used.

**Energy Distance:** Energy Distance measures the expected difference between pairs of samples from two distributions and provides a smooth, computationally efficient metric for comparing distributions:

$$E(\mathbf{X}_1, \mathbf{X}_2) = 2\mathbb{E}[\|\mathbf{X}_1 - \mathbf{X}_2\|] - \mathbb{E}[\|\mathbf{X}_1 - \mathbf{X}_1'\|] - \mathbb{E}[\|\mathbf{X}_2 - \mathbf{X}_2'\|],$$

where $\mathbf{X}_1$ and $\mathbf{X}_2$ represent matrices of vectors, and $\mathbf{X}_1'$ and $\mathbf{X}_2'$ are independent copies of $\mathbf{X}_1$ and $\mathbf{X}_2$. Energy Distance primarily matches the marginal characteristics of the distributions, such as mean and variance, without explicitly aligning their spatial structures. This makes it computationally efficient and ideal for quickly moving the optimization towards higher probability regions in the latent space due to its smoother loss surface (Székely & Rizzo, 2013).

**Sinkhorn Distance:** Sinkhorn Distance is a computationally efficient approximation of the Wasserstein distance (also known as Earth Mover's Distance), which measures the cost of optimally transporting one probability distribution to match another. The classic Wasserstein distance is defined as:

$$W(\mathbf{X}_1, \mathbf{X}_2) = \min_{\pi \in \Pi(\mu,\nu)} \sum_{i,j} \pi_{ij} c(\mathbf{X}_{1,i}, \mathbf{X}_{2,j}),$$

where $\Pi(\mu, \nu)$ represents the set of all transport plans between distributions $\mathbf{X}_1$ and $\mathbf{X}_2$, and $c(\mathbf{X}_{1,i}, \mathbf{X}_{2,j})$ is typically the squared Euclidean distance between vectors $\mathbf{X}_{1,i}$ and $\mathbf{X}_{2,j}$. The Wasserstein distance captures detailed structural characteristics such as covariance and spatial distribution of the data, making it particularly valuable for ecological applications (Villani, 2008). However, it is computationally expensive, especially in high-dimensional settings typical of environmental data. Sinkhorn Distance introduces an entropic regularization term to the Wasserstein distance, which not only smooths the optimization landscape but also makes the transport plan differentiable, allowing gradient-based optimization. The Sinkhorn distance is defined as:

$$S(\mathbf{X}_1, \mathbf{X}_2) = \min_{\pi \in \Pi(\mu,\nu)} \sum_{i,j} \pi_{ij} c(\mathbf{X}_{1,i}, \mathbf{X}_{2,j}) + \epsilon \sum_{i,j} \pi_{ij} \log(\pi_{ij}),$$

where $\epsilon$ is a regularization parameter that controls the trade-off between transport cost and entropy. The Sinkhorn algorithm iteratively adjusts dual variables to find a stable transport plan, which is computationally efficient and differentiable, unlike traditional optimal transport solutions. This differentiation enables the use of gradient descent for embedding optimization, making Sinkhorn particularly suitable for complex, high-dimensional comparisons (Cuturi, 2013).

#### Zero-shot Optimization Details

**Monte Carlo Sampling for Loss Estimation:** A key aspect of the optimization approach for zero-shot species is the Monte Carlo sampling of environmental vectors. At each optimization step, $N_{\text{pred}}$ samples of predicted environmental vectors, $\mathbf{E}_{\text{pred}}$, are drawn from the generative model conditioned on the current species embedding $\mathbf{z}_{s^*}$. This results in a matrix of predicted environmental vectors, $\mathbf{E}_{\text{pred}}$, which is compared to the matrix of observed environmental vectors $\mathbf{E}_{\text{true}}$. The flexibility of Energy and Sinkhorn distances allows them to operate on matrices with differing numbers of rows, meaning $N_{\text{pred}}$ does not have to match the number of observed points $N_{\text{true}}$. This flexibility is advantageous because it permits balancing between computational efficiency and optimization effectiveness. A larger $N_{\text{pred}}$ results in more accurate distance estimation but increases computational costs, while a smaller $N_{\text{pred}}$ adds stochasticity, which can help escape local minima during stochastic gradient descent. In practice, I found $N_{\text{pred}} = 1000$ samples per optimization iteration provided a good balance, offering sufficient stochasticity without excessively slowing down computation, particularly for test species with approximately 100 observed occurrence points. A good approach to the optimization was to have the parameter $\alpha$ start at 1 and gradually decrease it to 0 throughout the optimization. Initially, Energy Distance is emphasized due to its computational efficiency and smoother loss surface, which facilitates rapid convergence toward higher probability regions in the latent space. As $\alpha$ decreases, the focus shifts towards Sinkhorn Distance, which allows for fine-tuning by capturing intricate distributional details, albeit with a more complex loss landscape. The differentiability of both Energy Distance and Sinkhorn Distance is crucial, as it allows for efficient backpropagation of gradients through the loss function, enabling gradient-based optimization of $\mathbf{z}_{s^*}$.

### Model Architecture Details

#### NichEncoder

**Stage 1: Conditional Variational Autoencoder (CVAE)** The first stage of NichEncoder is a Conditional Variational Autoencoder (CVAE) that generates environmental variables $\mathbf{e}$ from species-specific embeddings, $\mathbf{z}_{\text{species}}$. The CVAE conditions on $\mathbf{z}_{\text{species}}$ by concatenating these embeddings to each layer of both the encoder and decoder networks, allowing the model to learn species-specific environmental niches. The architecture consists of multiple layers of Multi-Layer Perceptrons (MLPs) that encode and decode the input environmental variables.

##### Encoder and Decoder Structure

The encoder network takes environmental variables concatenated with species embeddings as input and processes them through three fully connected layers, each with 1024 neurons and ReLU activations. The output of the encoder includes the mean ($\mu$) and log variance ($\log \sigma^2$) of the latent variable $\mathbf{z}_{\text{VAE}}$. The decoder network mirrors this structure, taking the latent variable $\mathbf{z}_{\text{VAE}}$ and species embeddings as input to reconstruct the environmental variables $\hat{\mathbf{e}}$. Species embeddings are learned using an embedding layer that maps species identifiers to $\mathbf{z}_{\text{species}}$ vectors, which are used throughout the encoder and decoder networks.

##### Loss Function and Optimization Strategy

The CVAE is optimized using a combined loss function that includes the reconstruction loss, KL divergence, and species latent space regularization. The $\gamma$ parameter plays a crucial role in balancing the contributions of the likelihood and the KL divergence during training, progressively adjusting the importance of these terms to ensure the latent variables capture the data manifold effectively.

The total loss function is given by:

$$\mathcal{L} = \gamma \cdot \mathbb{E}_{q(\mathbf{z}_{\text{VAE}} | \mathbf{e}, \mathbf{z}_{\text{species}})}[\log p(\mathbf{e} \mid \mathbf{z}_{\text{VAE}}, \mathbf{z}_{\text{species}})] - D_{\text{KL}}(q(\mathbf{z}_{\text{VAE}} \mid \mathbf{e}, \mathbf{z}_{\text{species}}) \parallel p(\mathbf{z}_{\text{VAE}})) + \lambda \cdot \Omega_{\text{species}},$$

where $\gamma$ is initially set to down-weight the likelihood term and is gradually increased during training to enhance the model's focus on data reconstruction. The species regularization term, $\Omega_{\text{species}}$, is designed to keep the species latent space compact by penalizing a combination of squared (L2) and absolute (L1) values of the latent variables, effectively balancing between Gaussian (ridge regression) and Laplace (lasso regression) priors:

$$\Omega_{\text{species}} = (1 - \alpha) \cdot \sum (\mathbf{z}^2_{\text{species}}) + \alpha \cdot \sum |\mathbf{z}_{\text{species}}|,$$

where $\alpha$ is a tunable parameter controlling the weighting between the two regularization terms.

The optimizer used for training the CVAE is AdamW with a learning rate of 0.002, and a one-cycle learning rate policy is employed over 2500 epochs to dynamically adjust the learning rate during training.

**Stage 2: Rectified Flow Model** The second stage of NichEncoder uses a Rectified Flow model, inspired by (Liu, Gong, & Liu, 2022), to address the challenge of modeling the non-Gaussian posterior distribution of $\mathbf{z}_{\text{VAE}}$. This model estimates an Ordinary Differential Equation (ODE) that transforms noise into the complex distribution observed in $\mathbf{z}_{\text{VAE}}$.

##### Rectified Flow Model Details

The Rectified Flow model is based on a U-net style architecture adapted with MLP layers for vector field estimation (Figure 2, main manuscript). The architecture consists of three encoding and three decoding layers, with progressively decreasing and then increasing neuron counts (512, 256, and 128). Inputs to the model include the coordinates of latent variables, a time variable encoding, and species embeddings.

##### Training Data Creation

Training data for the Rectified Flow model is generated by interpolating between Gaussian noise samples and target samples from the latent distribution, forming a path that the model learns to approximate. The model is trained to predict the vector direction of these interpolated samples, effectively estimating a vector field pointing towards areas of high density in the target distribution.

##### Training Process

The training process of the Rectified Flow model consists of two stages:

1. In the first stage, the model learns to estimate the vector field that guides noise samples toward the target latent distribution. The model is trained on paths created by the interpolation, capturing the flow dynamics through the latent space.
2. In the second stage, the model refines the learned ODE by training on noise samples and their corresponding points after the initial ODE transformation. This rectification stage adjusts the ODE to approximate a linear transformation from noise to the target distribution, enhancing sampling efficiency.

##### Loss Function and Optimization Settings

The loss function minimizes the mean squared error (MSE) between the predicted and target vectors during training:

$$\mathcal{L} = \text{MSE}(\text{predicted vectors}, \text{target vectors}).$$

The optimizer used is AdamW with a learning rate of 0.001 and a weight decay of 0.01. A one-cycle learning rate policy is applied over 6000 epochs, adjusting learning rates dynamically to ensure effective training.

##### Trajectory Sampling

Trajectories are sampled using an ODE solver (`ode45`), integrating the learned vector field to transform noise into samples from the target distribution efficiently. This approach allows the model to generate high-quality samples with minimal integration steps.

#### GeODE

##### Model Architecture

GeODE uses a modified rectified flow architecture similar to that used in NichEncoder but tailored specifically for geographic data. The model generates 2-dimensional noise vectors as inputs, which are transformed through the rectified flow mechanism to output the desired geographic coordinates. The input consists of random noise vectors representing initial guesses in 2D space, while the conditioning input ($\mathbf{e}$) comprises environmental variables associated with each geographic location. Each environmental vector is normalized using means and standard deviations calculated from the data, ensuring numerical stability during training.

##### U-net Architecture

The core of GeODE is a U-net style structure implemented with Multi-Layer Perceptrons (MLPs) instead of convolutional layers . The U-net consists of two primary paths: downsampling and upsampling. In the downsampling path, the input noise vectors and environmental conditioning are passed through three fully connected layers with progressively smaller neuron counts (512, 256, and 128). These layers reduce the dimensionality while learning broad, high-level representations of the relationship between geographic locations and environmental factors. The upsampling path reconstructs the geographic coordinates by reversing the dimensionality reduction, using three corresponding fully connected layers to produce the final outputs. Skip connections between the downsampling and upsampling paths retain and propagate finer details, leading to more accurate predictions.

**Input Conditioning and Encoding**
In addition to the U-net structure, GeODE includes specialized encoding layers for the time variable $t$ and environmental conditioning vectors. A linear layer encodes the time step, representing the interpolation factor between noise and target coordinates. Another linear layer processes the environmental vectors, embedding them into a latent space that informs the transformation from noise to geographic coordinates. These encoded time and environmental vectors are concatenated with the latent representations from the U-net, allowing the model to incorporate both spatial and environmental dependencies into its predictions.

##### Training Data Creation

Training data for GeODE is generated through a Monte Carlo sampling process. Gaussian noise samples are drawn for both the latitude and longitude dimensions, creating initial random coordinate sets. These coordinates are linearly interpolated with target coordinates (actual occurrence points), guided by the ODE. This interpolation path forms the input for training, allowing the model to learn how to evolve from noise to realistic geographic distributions.

##### Training Process

**Stage 1: Vector Field Estimation**
In the first stage, the model learns to estimate a vector field that guides the initial noise samples toward the target coordinates, which represent actual geographic occurrence points. The U-net predicts the transformation vectors that align the noise vectors with the target distribution. Training data is generated by sampling Gaussian noise vectors for both $X$ and $Y$ dimensions, which are then linearly interpolated with the actual occurrence points. This interpolation forms a path between the noise and the target coordinates, which the model learns to follow.

**Stage 2: ODE Rectification**
The second stage refines the transformation by rectifying the ODE. In this step, the ODE is adjusted so that it evolves the noise vectors in a near-linear path toward the target coordinates, minimizing the computational steps required to generate realistic samples during inference.

##### Loss Function and Optimization

The loss function used to train GeODE is the mean squared error (MSE) between the predicted and target coordinates:

$$\mathcal{L} = \text{MSE}(\text{predicted coordinates}, \text{target coordinates}).$$

Optimization is performed using the AdamW optimizer with a learning rate of 0.001 and a weight decay of 0.01. A one-cycle learning rate policy is employed over 5000 epochs, dynamically adjusting the learning rate to improve training efficiency.

#### Latent Niche Space Interpolation

NicheFlow estimates a continuous latent representation of each species's niche, which exist in a latent niche space. From the vector a species's distribution in environmental space can be generated from the model. Indeed a hypothetical environmental distribution can be generated from any point in this latent niche space, includes latent vectors that do not correspond to any extant species in the dataset. To demonstrate this capability of the model, and to hopefully inspire readers to imagine potential uses of this capability, I provide an example in Figure 1 of this Supporting Information.

Figure 1: An example of interpolating between the latent niche vector of a reptile species from Northern South America to that of a reptile species from Southern South America. The top panel shows the latent vector as a bar chart. The two bottom left panels show four environmental variables (out of 32), showing generated distribution in pairs of two variables for the corresponding latent vector, based on the NichEncoder part of the model. The bottom right panel shows the generated distirbution in geographic space after passing the generated environmental data through GeODE. In this case it appears the interpolation causes the latent vector to pass through a part of niche space more characteristic of African environments than those of South America, including a substantial excursion in Mediterranean regions of South Africa and then around the Mediterranean sea. Interestingly, the west coast of South America which falls between the interpolated species is characterized by Mediterranean environments. It would be possible to restrict the output of the model to within South America but this shows successfully that the independence of the GeODE model from the species' geographic distributions allows generalized predictions of distributions to other continents and potentially future climates.

### Relationship of NicheFlow to traditional SDM methods

NicheFlow represents a new paradigm for Species Distribution Modelling which is fundamentally a joint method of SDM -- it relies on discovering patterns across many species for its strength. However, it is worth discussing how NicheFlow is related to other approaches.

#### NicheFlow fits a distribution in environmental space

NicheFlow conceptualizes a species' distribution primarily within environmental space, fitting a generative model to occurrence points in environmental space rather than geographic space using NichEncoder. It then uses GeODE to project the generated distribution of a species into geographic space. Importantly, GeODE is trained on environmental data and geographic coordinates without reference to any species occurrence points. This approach is most closely related to SDM methods that estimate probability density within environmental space, the most prominent example of which is probably the hypervolume approach of Blonder et al (). The hypervolume R package uses a method based on kernel density estimation to model the density of species occurrence points. Though the method was primarily designed to estimate niche volume and overlap of species, it also can be used to model species geographic distribution by projecting probabilities in environmental space to geographic space, though using a simpler method to do so than NicheFlow uses.

In essence hypervolume uses kernel density estimation to estimate $P(E|S)$ for each species of interest, where $E$ represents a set of environmental variable and $S$ is the occurrence of species $s$. This is then projected to geographic coordinates by assuming a set of grid points on the Earth are each associated with a unique environmental vector, generally defined as the environmental value of the mid-point of the grid-cell, let's call it $e_{x,y}$, and calculating the density of the species at point as $P(E = e_{x,y}|S)$. This can be related to NicheFlow by realizing that this procedure assumes a one to one correspondence between a coordinate and an environment, so that we can calculate $P(X, Y|E)$ simply as:

$$P(x_i, y_i|e_i) = 1, amp; \text{if } e_i = e_{x_i,y_i} 0, amp; e_i \neq e_{x_i,y_i}$$

Which essentially cancel the first term in the NicheFlow main text equation 1.

Note that hypervolume directly estimates this probability distribution whereas NichEncoder only learns how to generate from it.

The hypervolume approach, while appealing in its conceptual closeness to estimating the species environmental niche, similar to the philosophy of the NicheFlow model, has a number of limitations. A major one is computational efficiency. Kernel density estimation in general suffers strongly from the curse of dimensionality, where even a moderate number of dimensions can become quickly computationally prohibitive. As an example, fitting a hypervolume to a single species in our dataset with 800 occurrence points and 32 environmental dimensions took close to 12 hours on a regular working laptop computer with default setting. In order to estimate models for the ~10,000 species here would take 13 years without parallelization. More importantly however, is that kernel density methods, though excellent for estimating complex probability density distributions, are not easily extended to a multi-species setting. Though conditional kernel density estimation methods exist, methods that allow for conditioning variables to be latent, which is necessary to model generalizable variation between different species, are not well developed. In fact, methods based on generative deep learning provide the strongest promise is this area, and this is why NicheFlow uses these methods.

#### Most traditional SDM models fit a distribution in geographic space?

Most popular traditional SDM methods are considered to be fitting a model in geographic space because they don't explicitly estimate probability density in environmental space but rather attempt to distinguish geographic presence points from a set of non-presence points, using environmental variables as predictors. However, since explicit absence points are usually unavailable, non-presences are represented by a set of 'background' points drawn from a region of interest, usually randomly or sometimes on a grid. In this typical setting, presence and background points actually represent samples from two different environmental distributions: $P(E|S=s, O=1) = P(E|S)$, and $P(E|O=0) = P(E)$, where $O=1$ refers to the species being present (e.g. occurring), and $O=0$ is no species occurrence, e.g. non-presence or background.

For example the MaxEnt algorithm, when properly normalized, has been shown to estimate a quantity proportional to $P(E|S)/P(E)$, where $E$ represents environmental variables and $S$ represents species presence (Elith et al., 2011). This relationship arises from MaxEnt's equivalence to a Poisson point process model (Renner & Warton, 2013), where the intensity function $\lambda(E)$ is proportional to $P(E|S)/P(E)$.

#### Generalization to Presence-Background SDMs

However, this relationship is not unique to MaxEnt but extends more generally to Species Distribution Models (SDMs) that use random pseudo-absence or background points. We can formalize this connection as follows:

Let $Y$ be a binary variable where $Y=1$ for presence points and $Y=0$ for background points. In a presence-background classification, we estimate $P(Y=1|E)$. Applying Bayes' theorem:

$$P(Y=1|E) = \frac{P(E|Y=1)\cdot P(Y=1)}{P(E|Y=1)\cdot P(Y=1) + P(E|Y=0)\cdot P(Y=0)}$$

We can equate $P(E|Y=1)$ with $P(E|S)$, and $P(E|Y=0)$ with $P(E)$, assuming background points are drawn randomly from the available environment. If we assume $P(Y=1)$ is small, which is often the case for presence-background data, we can approximate:

$$P(Y=1|E) \approx c \cdot \frac{P(E|S)}{P(E)}$$

where $c$ is a constant.

To formalize this for a general presence-background SDM:

The model estimates a function $f(E)$ that maximizes the likelihood:

$$L = \prod_{i \in \text{pres}} f(E_i) \cdot \prod_{j \in \text{back}} (1 - f(E_j))$$

At the optimum, $f(E)$ will be proportional to $P(Y = 1|E)$. Given the approximation above, this means $f(E)$ is also proportional to $P(E|S)/P(E)$.

This formalization helps explain the often similar results produced by different SDM methods (e.g., MaxEnt, GLMs, GAMs) when applied to the same data. They are all estimating quantities proportional to $P(E|S)/P(E)$, albeit using different algorithms (Renner & Warton, 2013). It's important to note that this interpretation assumes that background points are drawn randomly from $P(E)$, and that presences are rare compared to the background. In practice, the constant of proportionality is often unknown, which is why SDM outputs are typically interpreted as relative rather than absolute measures of suitability. This relationship between presence-background classification and the estimation of $P(E|S)/P(E)$ provides a theoretical foundation for understanding traditional SDMs in the context of environmental space density estimation. It also offers a point of comparison for novel methods that more explicitly model in environmental space.

When applied to geographic space, traditional SDMs follow the same general procedure as hypervolume does, by assuming each grid-point in an area of Earth has a particular environmental vector associated with it $e_{x,y}$. The estimated function is then evaluated at each grid-point, e.g. $f(E = e_{x,y})$.

Unlike kernel density estimation methods, machine learning classification algorithms and statistical methods used to estimate $f(E)$ are fairly easy to extend to conditional models, and particularly for advanced hierarchical statistical models, latent conditioning variables are possible to incorporate. This allows modelling multiple species in a way that allows generalization because species can be represented in a latent vector space that can be estimated during model fitting. This is the basis of most current joint Species Distribution Models (jSDMs). In these model latent variables are linked to species presence probabilities using linear functions, such that they estimate a low-rank approximation of the covariance matrix for species occurrence probabilities. This linearity limits traditional jSDMs to modeling simple pairwise correlations amongst species. This is why estimates from jSDMs are often interpreted in term of pairwise species interactions, though it has been pointed out elsewhere that this interpretation is only viable under strict assumptions (Poggiato et al., 2021).

NicheFlow removes this linear limitation, allowing the relationship between species' latent niche vectors and their distribution to be highly non-linear. Whereas traditional jSDM can be considered as methods to approximate species pairwise covariances, NicheFlow can be interpreted as approximating a low-dimensional representation of the overall shape of niches in environmental space (a distribution of distributions). It is important to note for the interpretation of latent vectors and for downstream applications of them that this representation space is a non-linear non-Euclidean space -- representing a low-dimensional Riemannian manifold embedded within the ambient data space (Shao, Kumar, & Fletcher, 2017).

#### Summary of different methods

If we ignore for now geographic projections, we can summarize the connections between the types of SDMs in term of how they model suitability in environmental space, $S(E|S)$ thusly:

Traditional geographic SDMs like MaxEnt:

$S(E|S) \propto \frac{P(E|S)}{P(E)}$

Environmental SDMs like hypervolume and NicheFlow:

$S(E|S) \propto P(E|S)$

When we model suitability in environmental space like this we are often interested in more than just predicting a species' geographic distribution, but in estimating the species' environmental niche. For this purpose, the difference between different SDM methods ultimately is whether suitability is standardized by $P(E)$, which we can think of as the availability of different environments in the study region. There are distinct disadvantages of accounting for availability in this way for estimating a species' niche.

#### Disadvantages of standardizing by environmental availability

An important disadvantage of standardizing for environmental availability when estimating a species niche is that it induces a dependence on the background area of interest chosen when modelling the species, since availability of different environments may be different in different regions. Intuitively we consider the environmental niche of a species to be a property of the species, and so it should be constant with respect to the region under consideration. This ultimately makes traditional SDMs quite unsuitable to be used as estimates of the niche. This only allows relative comparisons of niches, for a fixed study area extent.

Another disadvantage is interpretation. Many researchers would naturally like to interpret suitability as proportion to the probability density of species occurrence, whereas suitability is instead a probability density ratio, which may be more difficult to interpret, especially without considering environmental availability. This could in some cases lead to results that are non-intuitive without proper consideration of environmental availability in the region of application.

A third disadvantage is what happens when you apply the model to environmental conditions that were not available in the background training data. If there was no data for that environment in the training data then the model is forced to extrapolate in these regions of environmental space, which can produce surprising or even nonsensical predictions that are highly sensitive to the assumption of the algorithm used, e.g. a GLM with extrapolate linearly outside the range of training data, in some case creating extremely unrealistic suitability values (Warren, Beaumont, Dinnage, & Baumgartner, 2019).

On the other hand, methods based on estimating $P(E|S)$ directly will also have no data where there is no availability of environments, however, extrapolation into these regions tends to at least produce less strange suitabilities because methods for estimating probability densities like kernel density estimation and generative models employ an assumption that areas with no data have a density of zero. This is generally a consequence of models that only use presence data, a lack of presence points is evidence for low density. For traditional methods, points that do not exist in the training data do not provide evidence one way or another, so that patterns from the data that do exist are just extrapolated to other parts of the space.

Methods like hypervolume and NicheFlow that model suitability as proportional to $P(E|S)$ do not have any of the above disadvantages, and they align well with the way we think about environmental niches. They have the added advantage that they do not require background points, removing the necessity to choose an often somewhat arbitrary background area of interest before beginning modelling. This also removes the need to extract large amounts of environmental data from typically large numbers of background points required for accurate estimates, and thus relieving a large computational speed and memory bottleneck in SDM.

However, modelling environmental suitability as directly proportional to probability density in environmental space, though aligning better with our conception of the environmental niche, does have one potential drawback. And that is the fact that environmental availability is not accounted for by the model in any way. In practice this means that $P(E|S)$ implicitly includes environmental availability as part of its estimated probability density. Another way of saying this is that the model cannot tell the different between an environment not existing in the species' suitability distribution, from the case where the environment simply does not exist at all. Generally speaking we believe that a species' niche could include environments that do not actually exist (perhaps due to evolution in past environments), or else not currently available to a species (perhaps due to dispersal limitation). The only way to possibly infer the existence of suitability beyond environments that actually exist in the presence data is with extrapolation of some kind. As discussed above, traditional methods could be used for this purpose if they produce reasonable extrapolation. In general this would likely require the embedding of realistic assumptions about how we expect niches to behave, as prior distributions in statistical models or as useful inductive biases in machine learning algorithms. However, most traditional SDMs use machine learning algorithms designed for general prediction whose assumptions are mostly idiosyncratic with respect to any species' biology.

A potential way around this drawback would be to incorporate environmental availability explicitly into the model as additional conditioning information, e.g. by creating a generative model for the probability distribution $P(E|S, A_e)$, where $A_e$ is some measure of the availability of an environment in an area of interest. This could be derived from an estimate of $P(E)$ itself or some set of covariates expected to covary with availability. A similar approach has been proposed for incorporating sampling bias into traditional SDMs (Warton, Renner, & Ramp, 2013), which is a related problem. Once the model is estimated, one could then generate from the model whilst fixing the environmental availability to some reasonable constant value as a way of removing the influence of availability from the model outputs.
